# Narrow window data-independent acquisition on the Orbitrap Astral Mass Spectrometer enables fast and deep coverage of the plasma glycoproteome

**DOI:** 10.1101/2024.07.29.605591

**Authors:** Shelley Jager, Martin Zeller, Anna Pashkova, Douwe Schulte, Eugen Damoc, Karli R. Reiding, Alexander A. Makarov, Albert J.R. Heck

## Abstract

Recently, a conceptually new mass analyzer was introduced by pairing a quadrupole Orbitrap mass spectrometer with an asymmetric track lossless (Astral^™^) analyzer. This system provides >200-Hz MS/MS scanning speed, high resolving power and sensitivity, and low-ppm mass accuracy. This instrument allows a narrow-window data-independent (nDIA) strategy, improving sensitivity and reproducibility even when using very short LC gradients. Although this represents a new technical milestone in peptide-centric proteomics, this new system has not yet been evaluated for the analyses of very complex and clinically important proteomes, such as represented by the plasma glycoproteome. Here, we evaluated the Orbitrap Astral mass spectrometer for the analysis of the plasma glycoproteome, and pioneer a dedicated nDIA workflow, themed nGlycoDIA. With substantially adjusted parameters and varying collision energies, nGlycoDIA has clear benefits for plasma glycoproteomics. We tested our method both in glycopeptide enriched and crude plasma, leading to the identification of more than 3000 unique glycoPSMs from 181 glycoproteins, covering a dynamic range of 7 orders of magnitude in the enriched plasma sample in just 40 minutes. In addition, we detect for the first time several glycosylated cytokines that have a reported plasma concentration in the ng/L range. Furthermore, shortening the gradient to 10 min still allows the detection of almost 2500 unique glycoPSMs from enriched plasma, indicating that high-throughput indepth clinical plasma glycoproteomics may be within reach.

## Introduction

Protein glycosylation is by far the most frequent and abundant co-/post-translational modification (PTM) of plasma proteins^1,2^. Compared to other PTMs, protein glycosylation is incredibly heterogenous, with many different possible glycan compositions and even more structural variations. Moreover, protein glycosylation is highly dynamic; glycosylation affects both biological properties of the protein, *e*.*g*., receptor interactions, immune response, and protein localization, as well as its physical properties, *e*.*g*., solubility, structure, and stability^3–6^. Glycan compositions and structures can change drastically upon external factors, such as diet, disease, and aging^2–4,7^. Recent advances in untargeted *N-*glycoproteomics, primarily performed using LC-MS/MS, have led to the identification of many putative glycan biomarkers from plasma and/or serum^8–13^. Although mass spectrometry-based glycoproteomics has been advancing at an incredible pace over the last decade, it is still facing major challenges compared to standard proteomics. Among these are signal dilution caused by glycan microheterogeneity, relatively poorer ionization efficiency of glycopeptides and the need for more complicated fragmentation methods to achieve glycan and peptide sequence information (i.e., stepping-HCD, EThcD). Plasma glycoproteomics is further complicated by the unfavorable high dynamic range of proteins occurring in plasma^14–17^. So far, to obtain a deep coverage of the glycoproteome, plasma glycoproteomics has generally been performed by analyzing enriched glycopeptides with LC-MS/MS using long LC gradients and data dependent acquisition (DDA).

In recent years, however, standard proteomics has been moving away from DDA and long LC gradients, shifting towards the higher throughput enabled by data independent acquisition (DIA). In DIA, instead of a single precursor, a broader mass window is chosen, leading to the selection of multiple precursors at once that are fragmented simultaneously. The resulting fragmentation spectra are then matched to either experimentally derived or theoretically generated spectral libraries^18^. While DIA is rapidly becoming the new standard for proteomics, it is not yet prevalent for glycoproteomics; software algorithms adopted for DIA data analysis, *e*.*g*., DIA-NN^19^ and Maxquant^20^, have not been developed or validated for glycopeptide library generation or annotation, although MSFragger^21^ recently added a library-based DIA-glycoproteomics workflow. Other developments in data analysis of DIA-derived glycopeptides include new algorithms for library generation (DIALib)^22^ or the practice of reformatting DIA into DDA type files, *e*.*g*., *via* DIA-Umpire, whereafter the data can be analyzed using a standard DDA data processing workflow^23–25^. Another recently reported strategy is to use DIA to screen for oxonium ion features, followed by DDA for further identification of the glycopeptides^26^. Lastly, Pradita *et al*. recently reported an untargeted DIA workflow on a purified protein, using Byonic for DIA-library generation and subsequent quantification in Skyline^27^. The latter still used manual curation for the matches generated by Skyline.

New advances in high-speed mass analyzers, such as the recently introduced Thermo Scientific™ Orbitrap™ Astral™ mass spectrometer (MS), push the field of standard peptide-based proteomics towards even more narrow windows for DIA, with MS^2^ isolation windows as small as 2 Th^28^. These small mass windows reduce the probability of chimeric spectra (resulting from the co-isolation of different precursors exhibiting alike *m/z*) and make the resulting data very similar to DDA, which typically uses similar precursor isolation windows. In addition, the very fast duty cycle of the Orbitrap Astral instrument, makes it possible to decrease the gradient length of the LC without sacrificing much depth in proteome coverage^29^. However, these powerful methods have not yet been fully tested and optimized for plasma *N*-glycoproteomics.

We here aimed to develop a strategy for DIA *N*-glycoproteomics on the Orbitrap Astral MS. As a benchmark to compare we used our current HCD-based workflow for plasma glycoproteomics, including an optional enrichment of glycopeptides using cotton-HILIC^30^, followed by LC-MS/MS utilizing stepping HCD^31^ and data analysis using Byonic^32^. With the here developed DIA *N-* glycoproteomics workflow we aim to optimally use the speed offered by the Orbitrap Astral MS, using narrow-window DIA to allow application of conventional DDA data analysis workflows. For this purpose, glycopeptides were enriched from human pooled plasma using cotton-HILIC SPE and analyzed using different gradient lengths and collision energies. Following this experimental setup, we observed that DIA on the Orbitrap Astral MS needed to be substantially optimized for the analysis of glycopeptides, primarily because of their generally higher masses and lower charges, resulting in higher *m/z* values, and their distinct fragmentation characteristics. We here demonstrate how our optimizations led to an unprecedented coverage of the plasma glycoproteome, even when using relatively short gradients lengths of 10 to 20 min, where identifications of glycoproteins span about 6 to 7 orders of magnitude in plasma concentrations. For comparison, when using crude plasma, without any enrichment or depletion, we can, using the same nGlycoDIA approach, cover glycosylation of proteins spanning 3 orders of magnitude. We foresee that such glycoproteomics directed methods would enable to measure plasma glycoproteome profiles in a 20 min timeframe per sample, using only 10 µL of plasma per donor, substantially increasing the throughput and depth of these clinically relevant analyses.

## Methods

### Sample preparation

10 µL of pooled plasma sample (VisuCon-F Normal Donor Set (EFNCP0125), Affinity Biologicals) was mixed with 40 volumes of SDC-buffer (0.1 M TRIS/HCl pH 8, 40 mM TCEP, 100 mM CAA, 1% SDC, *v/v*) for denaturation, reduction and alkylation. These samples were heated to 80 °C for 10 min, after which the proteases Trypsin and LysC were added in a 1:70 and 1:50 enzyme to protein ratio, respectively. Following overnight digestion at 37 °C, the samples were quenched with 0.1% TFA. The peptide digests were dried *in vacuo* and reconstituted in 0.1% TFA. Then, samples were loaded onto Oasis PRiME HLB (10 mg) 96-wells plates (Waters), washed with 0.1% TFA, and peptides were eluted using 60% ACN/0.1% TFA. The obtained digests were either analyzed directly by LC-MS/MS as the crude plasma sample or enriched for glycopeptides.

Glycopeptide enrichment was performed using cotton-HILIC SPE^33^. 100% cotton thread was cut into 10 mm lengths and individual threads were separated. Each thread was placed in a pipette tip, of which 8 at the time were used on a multichannel pipette. The tips were conditioned in 0.1% TFA (3x 100 µL) and washed with 80% ACN/0.1% TFA (3x 100 µL). Plasma peptide samples were reconstituted in 80% ACN/0.1% TFA (100 µL) and loaded onto the tips by 30 times repetitive pipetting. Tips were washed by three times repetitive pipetting in 80% ACN/0.1% TFA (3x 100 µL), and finally eluted by three times repetitive pipetting in 50% ACN/0.1% TFA (3x 100 µL). The three elution fractions were combined and dried *in vacuo*. Glycopeptide enriched samples were reconstituted in 100 µL 0.1% TFA and crude plasma samples were reconstituted in 1 mL 0.1% TFA for an approximately 200 ng/µL concentration. Peptide solutions were split in 12 µL batches in glass vials for LC-MS/MS analysis and 1 µL was injected per analysis.

### LC-MS/MS Analysis

Exploratory DDA data was acquired using a Thermo Scientific™ Ultimate™ 3000 HPLC system (Thermo Fisher Scientific, Germering, Germany) coupled online to a Thermo Scientific Exploris™ 480 mass spectrometer (Thermo Fisher Scientific, Bremen, Germany). Peptides were trapped using a Thermo Scientific PepMap Neo Trap Cartridge (5 μm C18 300 μm x 5 mm) separated on a 50 cm reverse phase analytical column packed in-house with integrated emitter (ReproSil-Pur 120 C18-AQ, 1.9 µm particles, 75 µm x 50 cm; Dr. Maisch). The trapping column was heated at 40 °C, and the analytical column was heated at 50 °C. The LC mobile phases used were water with 0.1% formic acid (solvent A) and 80% acetonitrile in water with 0.1% formic acid (solvent B) (both UPLC Grade, Biosolve). The column was equilibrated at 3% B, and kept there after injection for 1 min, then linearly increased to 13% B in 4 minutes, linearly increased to 44% B over 113 minutes, ramped up to 99% B in 2 minutes, kept constant for 4 minutes before decreasing to 3% B in 1 minute, followed by 10 minutes of equilibration. The flowrate was kept constant at 0.3 µL/min.

The Exploris 480 mass spectrometer was operated in positive ion mode, with a spray voltage of 2000 V and an ion transfer tube temperature of 275 °C. The MS^1^ settings were as follows: Orbitrap resolution 120,000 (FWHM at *m/z* 200); scan range *m/z* 350-2000; maximum injection time 50 ms. MS^2^ scans were recorded in DDA mode with a cycle time of 3 seconds, precursors were sorted in two steps: highest charge state first, followed by lowest *m/z*, and dynamic exclusion was set to 15 s. The MS^2^ settings were as follows: isolation window: 1.6 Th, NCE 20 and 40 % (stepped), Orbitrap resolution: 60,000, scan range: 120-4000 *m/z*, maximum injection time mode: auto.

All DIA data were acquired using a Thermo Scientific Vanquish™ Neo UHPLC (Thermo Fisher Scientific, Germering, Germany) coupled online to an Orbitrap Astral MS (Thermo Fisher Scientific, Bremen, Germany). The method order was randomized to reduce dilution effects, with the only condition that all enriched plasma samples were analyzed first to limit potential carry-over from the crude plasma samples. Peptides were separated on an IonOpticks Aurora Ultimate TS nanoflow UHPLC column (25 cm x 75 µm inner diameter, C18 stationary phase, 120 Å pore size, 1.7 µm particle size, IonOpticks PtyLtd, Fitzroy, Australia), in direct-injection configuration. The column was heated at 55 °C. The LC mobile phases used were water with 0.1% formic acid (solvent A) and 80% acetonitrile in water with 0.1% formic acid (solvent B) (both Optima LC/MS Grade, Fisher Chemical). For the DIA analysis, four different gradients were used: the concentration of solvent B was gradually increased from 1 to 4% B at a flowrate of 0.55 µL/min, to 8% B in 0.9 minute at a reduced flowrate of 0.3 µL/min, and to 30%, either in 10, 20, 30 or 40 minute, for the 10, 20, 30, and 40 minute gradient runs, respectively. Lastly, the column was washed by increasing solvent B to 99% over 3 minutes and maintaining this concentration for 3 minutes, both at a flowrate of 0.55 µL/min.

The Orbitrap Astral mass spectrometer was operated in positive ion mode, with a spray voltage of 2000 V and an ion transfer tube temperature of 290 °C. Before submission of the sequence, the mass spectrometer was calibrated in positive ion mode for FTMS mass accuracy, Astral mass accuracy as well as ion foil and prism 2 for ion transmission. For the standard DIA method, the MS^1^ settings were as follows: Orbitrap resolution 120,000 (FWHM at *m/z* 200); scan range *m/z* 380-980; maximum injection time 100 ms; AGC target 500%. The DIA parameters were as follows: isolation window: 2 Th; HCD collision energy: 27%; precursor mass range: *m/z* 380-980; scan range: *m/z* 150-2000; maximum injection time: 3 ms; AGC target: 200%. For the nGlycoDIA method, the MS^1^ settings were as follows: Orbitrap resolution 120,000 (FWHM at *m/z* 200); scan range *m/z* 950-1655; maximum injection time 100 ms; AGC target 500%. The DIA parameters were as follows: isolation window: 3 Th; HCD collision energy: 25%, 27%, 30%, or 35%; precursor mass range: *m/z* 950-1655; scan range: *m/z* 150-2000; maximum injection time: 4 ms; AGC target: 800%. For all DIA experiments a default charge state of two plus was used, and the DIA window placement optimization was used. The major experimental parameters used are summarized in **Table 1**.

**Table 1:** Overview and comparison of key experimental parameters in the DDA, standard nDIA and nGlycoDIA methods. Compared to the standard nDIA method, the optimized nGlycoDIA method differs primarily in the chosen MS^1^ scan range. A slightly larger sliding window was chosen to accommodate also for this higher m/z range and the broader isotope distribution of the larger glycopeptides. Excluding the most abundant “high” m/z ins originating from non-modified peptides (e.g. originating from albumin) we also could extend the AGC target and injection time.

|  | DDA method | Standard nDIA method | nGlycoDIA |
| --- | --- | --- | --- |
| **Orbitrap Resolution** | 240000 | 120000 | 120000 |
| **MS<sup>1</sup> Scan Range (m/z)** | 380-2000 | 380-980 | 950-1655 |
| **MS<sup>2</sup> Isolation window (Th)** | 1.6 | 2 | 3 |
| **MS<sup>2</sup> Maximum injection time (ms)** | 10 | 3 | 4 |
| **MS<sup>2</sup> AGC target (%)** | 100 | 200 | 800 |

### Data analysis

Raw data files were searched using PMI-Byonic (v.5.5.2, Protein Metrics). The protein database was a focused database consisting of plasma proteins (annotated to be in blood by using the Human Protein Atlas depository^34,35^) and common isoforms (with higher frequency than 5%) as can be found in NextProt^36^ (3107 entries), reverse decoys were added by Byonic. The rationale not to search against the entire human proteome database was to decrease search time. Fully specific digestion with at maximum two missed cleavages was allowed, and the precursor and fragment mass tolerance were set to 10 and 20 ppm, respectively. Carbamidomethylation of C was set as fixed modification, variable modifications included oxidation on M or W and pyroglutamic acid formation on protein and peptide N-terminal Q and E. A glycan list consisting of 279 glycans commonly observed in human proteins was used, as provided in the Supplemental information (**Supplemental Table 1**). In this table and in the text glycan compositions are abbreviated, where N or HexNAc is *N*-acetylhexosamine, H or Hex is hexose, F or dHex is fucose, P or Phos is phosphomannose, and S or NeuAc is *N*-acetylneuraminic acid/sialic acid. Each peptide was allowed a maximum of one variable modification and one glycan. Precursor and charge assignments were computed from the MS^1^ data and a maximum of two precursors was allowed per MS2 scan.

Subsequent data analysis was performed using R (v4.3.1). FDR filtering was based on the individual PSM scores: PSMs were ordered from high to low score and a cut-off was calculated per file to have no more than 1% reverse sequences. Then, data was filtered on the criteria that the PSM had to be identified in at least two out of four replicates within any of the measurement conditions. The package ComplexHeatmap was used for heatmaps and clustering. Ion coverage calculations were performed with an in-house developed script using Rust and rustyms (https://github.com/snijderlab/rustyms). All code will be available as **supplemental files S10 and S11**, for the R and Rust scripts, respectively.

### Structural glycan modeling

Structural modeling of the most abundant glycan on the cytokines IL-12, IL-22, and VIP was done using GLYCAM-Web^37^. Because our methods only yield compositional and limited structural information, but not linkage information, we modeled one specific structure per glycan. These were: NeuAc-α2,6-Gal-β1,4-GlcNAc-β1,2-Man-α1,3-(Man-α1,6-(Man-α1,3)-Man-α1,6-)Man-β1,4-GlcNAc-β1,4-GlcNAc-β1-Asn for N3H6S1, NeuAc-α2,6-Gal-β1,4-GlcNAc-β1,2-Man-α1,3-(Gal-β1,4-GlcNAc-β1,2-Man-α1,6-) Man-β1,4-GlcNAc-β1,4-GlcNAc-β1-Asn for N4H5S1, NeuAc-α2,6-Gal-β1,4-GlcNAc-β1,2-Man-α1,6(NeuAc-α2,6-Gal-β1,4-GlcNAc-β1,4(NeuAc-α2,6-Galβ1,4-GlcNAc-β1,2-)Man-α1,3-)Man-β1,4-GlcNAc-β1,4-GlcNAc-β1-Asn for N5H6S3

## Results

### Setting up narrow-window nGlycoDIA

To set up a glycoproteomics-directed narrow-window DIA method, *i*.*e*., nGlycoDIA, several experimental parameters needed to be considered and optimized, including *m/z* range, isolation window size, and maximum ion injection time. In DDA-mode, we typically use an MS^1^ scan range between *m/z* 380-2000 ^30,31,38,39^, however this is not ideal in the DIA method because it would increase the cycle time too much. To choose an efficient MS^1^ scan range, we examined data derived from glycopeptide enriched plasma analyzed on an Orbitrap Exploris MS. Here we noticed that glycopeptides typically have a substantially higher *m/z* distribution than unmodified peptides, in line with what has been reported earlier^13,31,40^, and likely due to their on average larger size/mass and reduced charge (**Supplemental Figure S1**). Based on this observed precursor ion distribution, we chose our glycoproteomics-specific DIA method to span an *m/z* range of 955-1650 (median ∼1300), which is quite distinct from, and nearly does not even overlap with, the standard proteomics range that has typically been used in DIA proteomics, *i*.*e*., *m/z* 380-980 (median ∼680)^28,29^. An additional advantage of this change in selected precursor *m/z* range, is that it effectively leads to a second degree of glycopeptide enrichment in the gas-phase. In our optimized nGlycoDIA method, the isolation window size was set to 3 Th (to reflect the higher median *m/z*) and the maximum injection time to 4 ms. (**Table 1**). A downside of shifting the MS^1^ range in nGlycoDIA, is that it leads to a 9-fold decrease in number of PSMs and a 4-fold decrease in unique PSMs for unmodified peptides in the crude plasma, reflecting that many more unmodified highly abundant tryptic peptides fall in the 380-980 window when compared to the 950-1655 window (**Supplemental Figure S2**). Fortuitously, with less peptides present, choosing the nGlycoDIA window did also lead to a substantial decrease in chimeric spectra (∼11% *versus* 4% of the matched spectra were chimeric for standard nDIA and nGlycoDIA, respectively), even although the isolation window in nGlycoDIA is broader. We chose not to increase the window further to minimize the potential increased occurrence of chimeric spectra. Furthermore, the AGC target was increased to 800% to fully use the 4 ms injection time.

### nGlycoDIA enables deep coverage of low abundant plasma proteins

Next, using the experimental parameters described above, a set of DIA methods were designed for the Orbitrap Astral MS exploring four different NCEs, namely 25, 27, 30, and 35%. Because DIA often results in identification of low abundant proteins and increased sensitivity^41,42^, we wanted to explore how many glycoproteins could already be observed without glycopeptide enrichment. Therefore, we analyzed in parallel both plasma samples from which the glycopeptides had been enriched by cotton-HILIC, as well as what we term here “crude” plasma samples (**Figure 1**). Additionally, as shown for standard proteomics with the introduction of 40 up to 180 samples per day (SPD) methods^28^, we hypothesized that the increased speed of the Orbitrap Astral MS would allow us to shorten the LC-MS runs, without a significant loss in identifications. We therefore evaluated four different LC gradients, going down from 40, 30, 20, to even 10 minutes.

**Figure 1:**
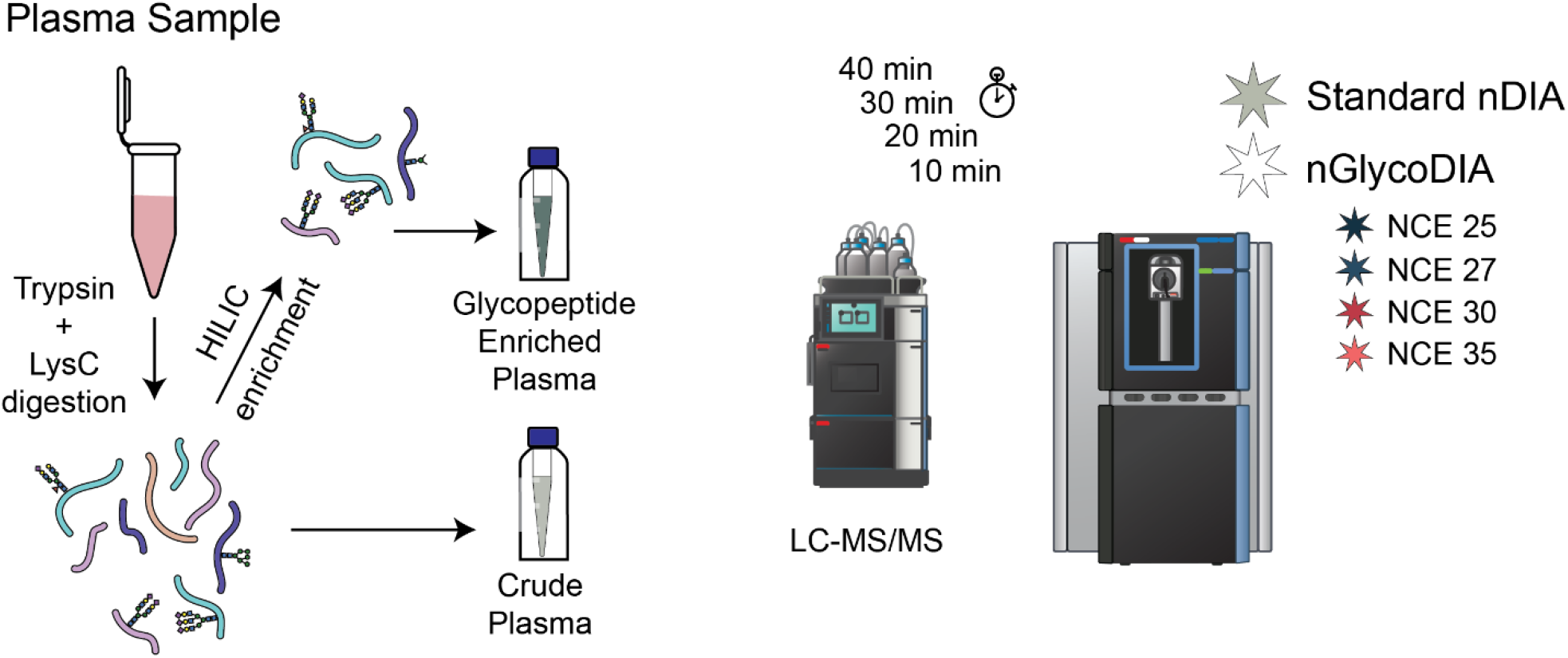
Evaluation of data-independent acquisition (DIA) of plasma glycopeptides. Plasma samples were proteolytically digested using trypsin and LysC. The resulting sample (crude plasma digest) was either directly subjected to LC-MS/MS analysis, or first enriched for glycopeptides by cotton-based hydrophilic interaction chromatography (HILIC). LC-MS/MS analysis was performed with different gradient lengths (10, 20, 30 and 40 min). Employed MS methods included both the standard DIA method, as typically used for non-modified peptides, as well as various optimized nGlycoDIA methods using different NCEs (25, 27, 30 and 35%).

First, the 40-minute gradient results were examined because we expected to identify the highest number of glycopeptides using this longest gradient. Higher fragmentation energy led to the identification of more unique glycopeptide PSM (glycoPSMs), their number steadily increasing across NCEs 25-35% (**Figure 2A**). This increase is attributed to both an increase in identified proteins and glycosites (up to NCE 30%), as well as an expansion in detected glycan microheterogeneity per site (**Figure 2B, C**). These NCE-dependent features were observed in measurements for both the crude and enriched plasma samples. Overall, the glycopeptide enrichment led to a substantial increase in unique glycopeptide identifications, compared to crude plasma. In the standard DIA method, a significantly lower number of glycopeptides was identified, most detections being non-glycosylated peptides instead. This is also represented in the number of glycoproteins and glycosylation sites. Interestingly, the increase in the number of co-enriched non-glycosylated peptides lead to an increase in non-glycosylated PSMs compared to the crude plasma, which is also represented in the number of identified proteins in each sample (252 in the enriched plasma compared to 184 in the crude plasma). Injection replicates were highly reproducible, with approximately 80% of identifications found in at least two replicates (**Supplemental Figure S3A and S3B**). Furthermore, most glycopeptides that were identified in two analyses were found in all four of the replicates (**Supplemental Figure S3A**). When comparing between NCEs, the overlap between the smaller datasets (acquired with lower NCE) was similar to the overlap between injection replicates, indicating that the increase in annotatable spectra was an increment and not a bias (**Supplemental Figure S3B**).

**Figure 2:**
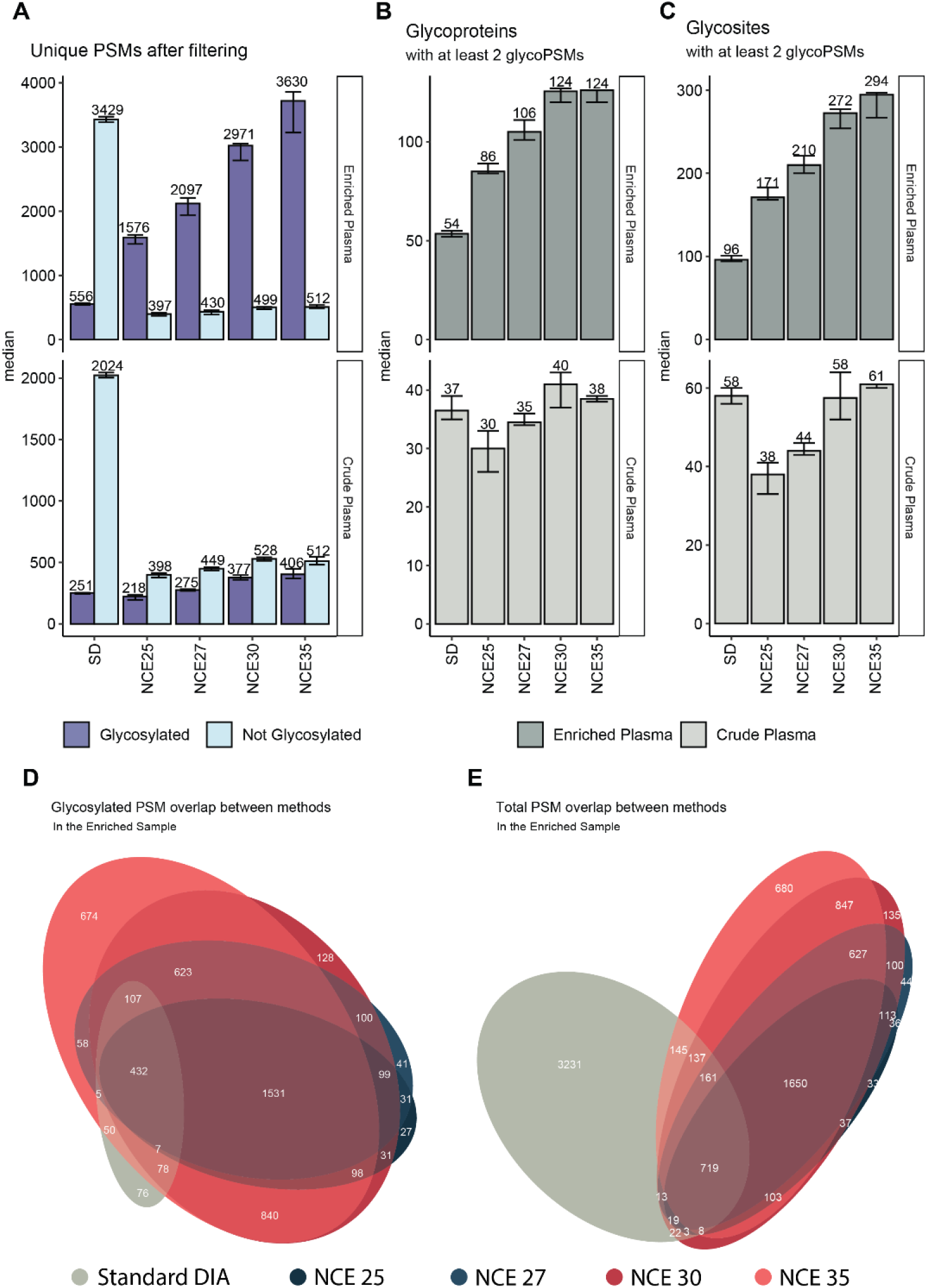
nGlycoDIA glycoproteomics data obtained for crude and glycopeptide-enriched plasma samples. The top three panels depict data for glycopeptide-enriched plasma, whereas the middle three panels depict data for crude plasma. **A**) The number of unique PSMs for glycopeptides (purple) and non-modified peptides (light blue). **B**) The number of proteins identified. **C**) The number of glycosites identified. The consecutive bars reflect the results obtained with the different DIA methods and NCE energies, as annotated. Four technical replicates were performed for each method. The bars illustrate the median values, while the error bars extend from the minimum to the maximum number identified out of the four replicates. The x-axis annotation on **A-C** refer to the MS/MS methods, where SD is standard DIA, and NCE25, NCE27, NCE30, and NCE35 refer to nGlycoDIA with the respective NCE percentages. **D**) A Venn-diagram of the PSM overlap between methods for the detected glycosylated PSMs. **E**) A Venn-diagram of the overlap of PSMs between methods considering all PSMs both glycopeptides and non-modified peptides. **D** and **E** are also available as upset plots in **Supplemental Figure** S**4A** and **S4B**. All data (**A-E**) was filtered for 1% FDR and, to be included, each unique PSM had to be identified in at least two out of four replicates for a single condition.

Next, the identifications extracted from the analysis of the crude plasma were compared to those identified in the glycopeptide-enriched plasma. Here we see that approximately 85% of the glycopeptide identifications in crude plasma were also found after enrichment. Overall, a high overlap (only 1-4% unique glycoPSMs per method, excluding NCE 35%) of identified PSMs between the different nGlycoDIA methods was observed for both glycosylated and unmodified peptides, indicating high method robustness (**Figure 3D and 3E, Supplemental Figure S4A and S4B**). Almost all identified glycosylated PSMs found in the standard DIA method were detected in each of the nGlycoDIA methods as well. This illustrates that the data lost by shifting the precursor *m/z* range is minimal, and that this rather results in a beneficial additional degree of enrichment in the gas phase.

**Figure 3:**
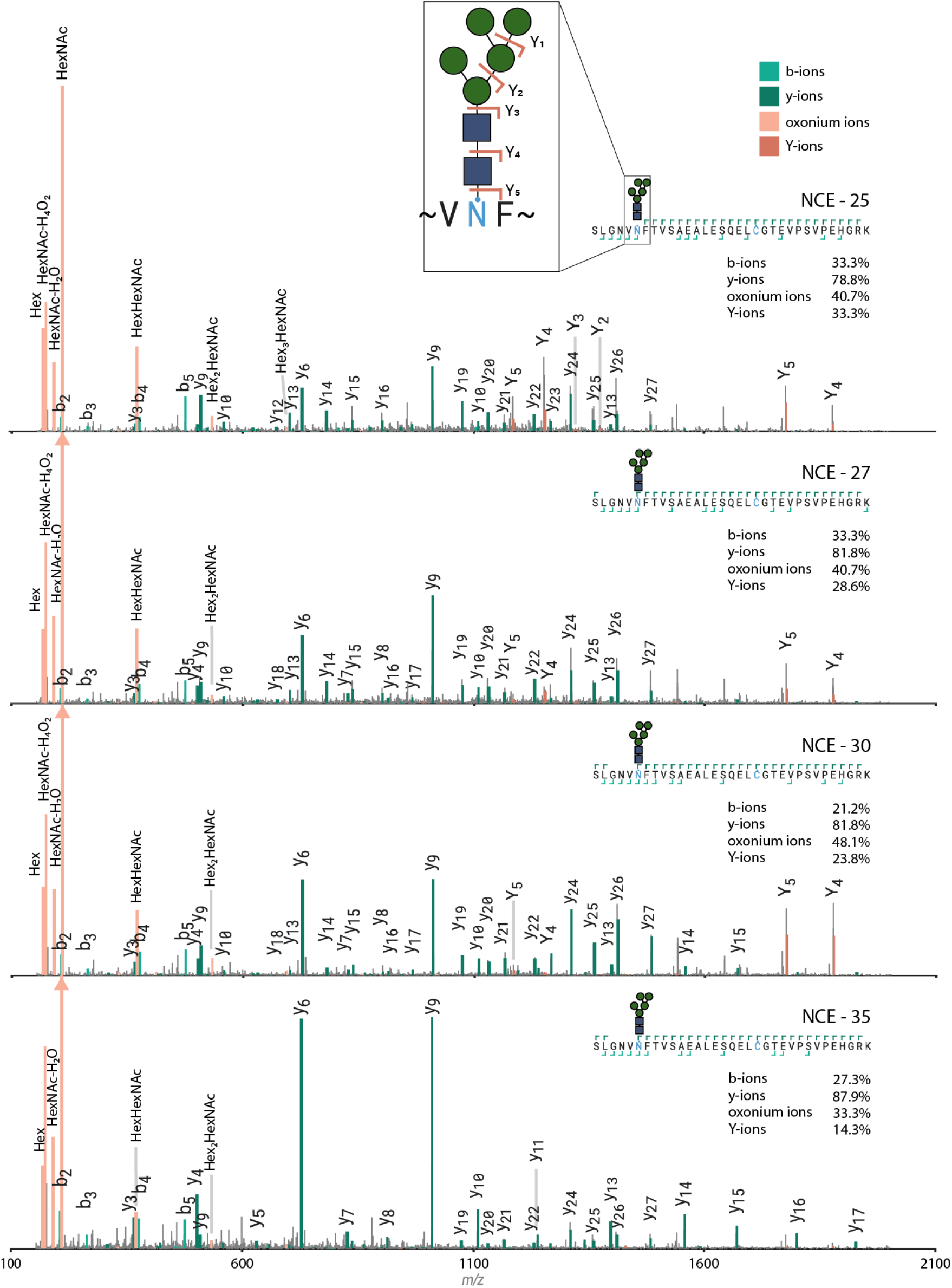
Effect of collision energy on glycopeptide fragmentation spectra. The highest scoring PSM for each specific peptide with glycan was selected per condition to ensure a fair comparison. In the figure, b- and y-ions are represented in light and dark green, respectively, while B-(oxonium) and Y-ions are depicted in light and dark orange, respectively. In the top right corners, the coverage of the peptide backbone is shown, with percentages of total possible positions covered by the respective ion. Annotations were performed using Annotator (v 0.2.2), which is accessible at https://github.com/snijderlab/annotator.

While the number of uniquely identified glycoPSMs kept increasing with higher collision energies, we observed that the quality of the MS^2^ spectra decreased from 25% NCE to 35% NCE. This especially became evident once examining the glycan specific ion coverage over the entire dataset (**Supplemental Figure S3C**). Both B-(oxonium) and Y-ion coverage decreased consistently over different precursor charge states. **Figure 3** serves as an example, demonstrating that by using lower collision energy, larger glycan fragments, as well as abundant Y-ions were preserved, which increased the confidence in both glycan and peptide assignment. These Y-ions are crucial when modifications of the peptide backbone might overlap in mass with different glycoforms. For instance, the mass difference between a hexose and a fucose is identical to that of a peptide with and without oxidation, *i*.*e*., 15.99 Da, the mass of an oxygen. Additionally, while Y-ions are precursor-specific, oxonium ions are not. As such, Y-ions are critical in determining glycan identity in the case of chimeric spectra.

Next, we used the blood protein concentrations reported in the Human Protein Atlas (which we will refer to as the Blood Atlas^34,35^) to explore the depth of proteome coverage we could reach using nGlycoDIA. In the crude plasma, primarily highly abundant glycoproteins (concentration > 1 mg/mL) were identified (**Figure 4A, B**). When comparing standard DIA to our optimized nGlycoDIA method, it was clear that in the latter not only more glycoproteins in the mg/L range were identified (51 with nGlycoDIA compared to 42 with standard DIA), but even lower-abundant glycoproteins such as Factor VIII and the epidermal growth factor receptor. Additionally, the glycan microheterogeneity that could be identified was overall higher when using nGlycoDIA (**Figure 4D)**. The glycans characteristics observed in the standard DIA method were also quite distinct and shifted towards lower mass glycans when compared to the nGlycoDIA methods. For example, in standard DIA we observed more N4H5S1 than N4H5S2, relatively more paucimannose glycans, and almost no tri- and tetra-antennary glycans (**Supplemental Figure S5**). This was not surprising, because the DDA experiments already demonstrated that most glycopeptides fall outside of the precursor range used in the standard DIA method (**Supplemental Figure S1)**. It should be noted that the method presented here focused on plasma *N-*glycoproteomics, where large complex-type glycans are highly abundant. When analyzing different types of samples where smaller glycans are more prevalent, such as paucimannose glycans, the MS^1^ scan range might require additional optimization. In all, there was no clear difference in glycan distribution between the different NCE conditions used, and the distribution of different glycans was very similar to other plasma glycomics and glycoproteomics studies, with glycopeptides bearing mostly sialylated complex type glycans with two antennae, followed by complex tri and tetra-antennary glycans^6,13,31,43^.

**Figure 4:**
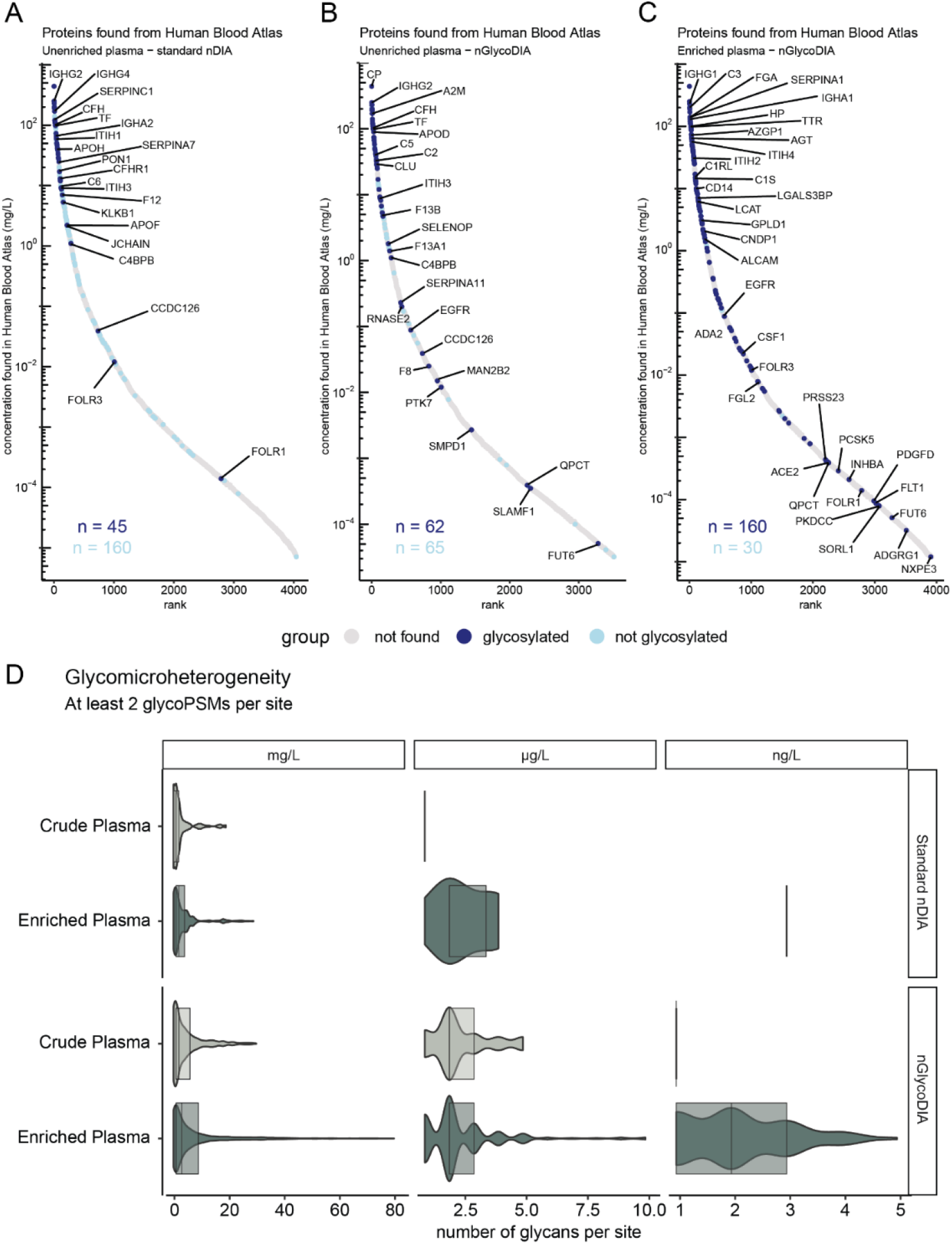
Evaluation of the depth and annotatable microheterogeneity of both high abundant and low abundant plasma glycoproteins following analysis by nGlycoDIA. **A**) the proteins detected in crude plasma using the DIA method optimized for standard proteomics. **B**) Protein detections when analyzing crude plasma digest by nGlycoDIA. **C**) Protein detections following glycopeptide enrichment and subsequent analysis with nGlycoDIA. All detected proteins were compared to their reported abundance in “normal plasma” as described for the blood proteins of the Human Protein Atlas (Blood Atlas), as measured by MS^35,53^. In the bottom-left corner, the number of proteins detected with and without identified glycosylated peptides is indicated in dark blue and light blue, respectively. **D**) Violin plot of number of unique glycans identified per site. Data is grouped on the reported abundance of the proteins in the Blood Atlas, where mg/L ranges between 999 - 1 mg/L, µg/L between 999 - 1 µg/L, and ng/L below 999 ng/L. The overlaying boxplot show the median value, and the values of the upper and lower quartile. The comparison revealed that nGlycoDIA in glycopeptide-enriched plasma led to most glycans detected per site, allowing the detection of glycoproteins across a dynamic range of 10^6^.

Following HILIC enrichment, substantially more glycoproteins were identified and consequentially a larger dynamic range was covered (spanning 7 orders of magnitude). Compared to the analysis of crude plasma with nGlycoDIA, the number of glycoproteins detected in the enriched sample in the mg/mL range was doubled, in the µg/mL range the number of identifications increased by over 4-fold and we were even able to identify 17 glycoproteins in the ng/mL range (**Figure 4C**). Additionally, 12 glycoproteins were identified that only have a reported quantification in the Blood Atlas by immunofluorescence and not by mass spectrometry. Moreover, 25 glycoproteins were detected that have not been quantified in the Blood Atlas at all, amongst them being IGHD, ITIH5, and intriguingly even several cytokines which we will discuss in more detail in the following section. As these proteins are currently not quantified in the human blood atlas, they cannot be shown in the S-curves in Figure 4. Next to increased detection of low-abundant proteins and glycosylation sites, for the glyco-enriched plasma we were able to detect more glycan structures per site compared to its crude counterpart. In the most extreme case, we observed more than 80 different glycans on a single site, namely N639 of serotransferrin, highlighting the visibility of glycosylation heterogeneity (**Figure 4D**). Overall, the data demonstrate that enrichment for glycopeptides is still highly beneficial for the analysis of low abundant glycoproteins (concentration < 1 mg/mL) in a complex matrix such as plasma.

### Benchmarking plasma glycoproteomics *versus* proteomics, detection of glycosylated low abundant cytokines

In the present study we identified 181 glycoproteins and 436 unique glycosites with the optimized nGlycoDIA approach, taking as a cut-off their detection in at least two out of four individual experiments (acquired at different NCE). We were intrigued by the detection of several glycosylated cytokines and growth factors in our study, whereby we here focus on and discuss only the ones that we detected in at least two experiments. These include the interleukins IL-12A, IL-12B and IL-22, the colony stimulating factors CSF1 and CSF2, and the vasoactive intestinal peptide (VIP). All these cytokines are known to play an eminent role in inflammation and diseases such as cancer^44–47^, but experimental evidence for their glycosylation in human plasma has as far as we know never been reported. The here provided insight into their glycosylation can provide a valuable background for elucidating the role of these PTMs in the disease mechanisms in which these cytokines play a role. Below we will describe in more detail the glycosylation features that we identified on these cytokines in the pooled healthy plasma (**Figure 6A**). Furthermore, to demonstrate the relevance of these glycans, we modeled the most abundant ones on structural models of the proteins, when available.

**Figure 5:**
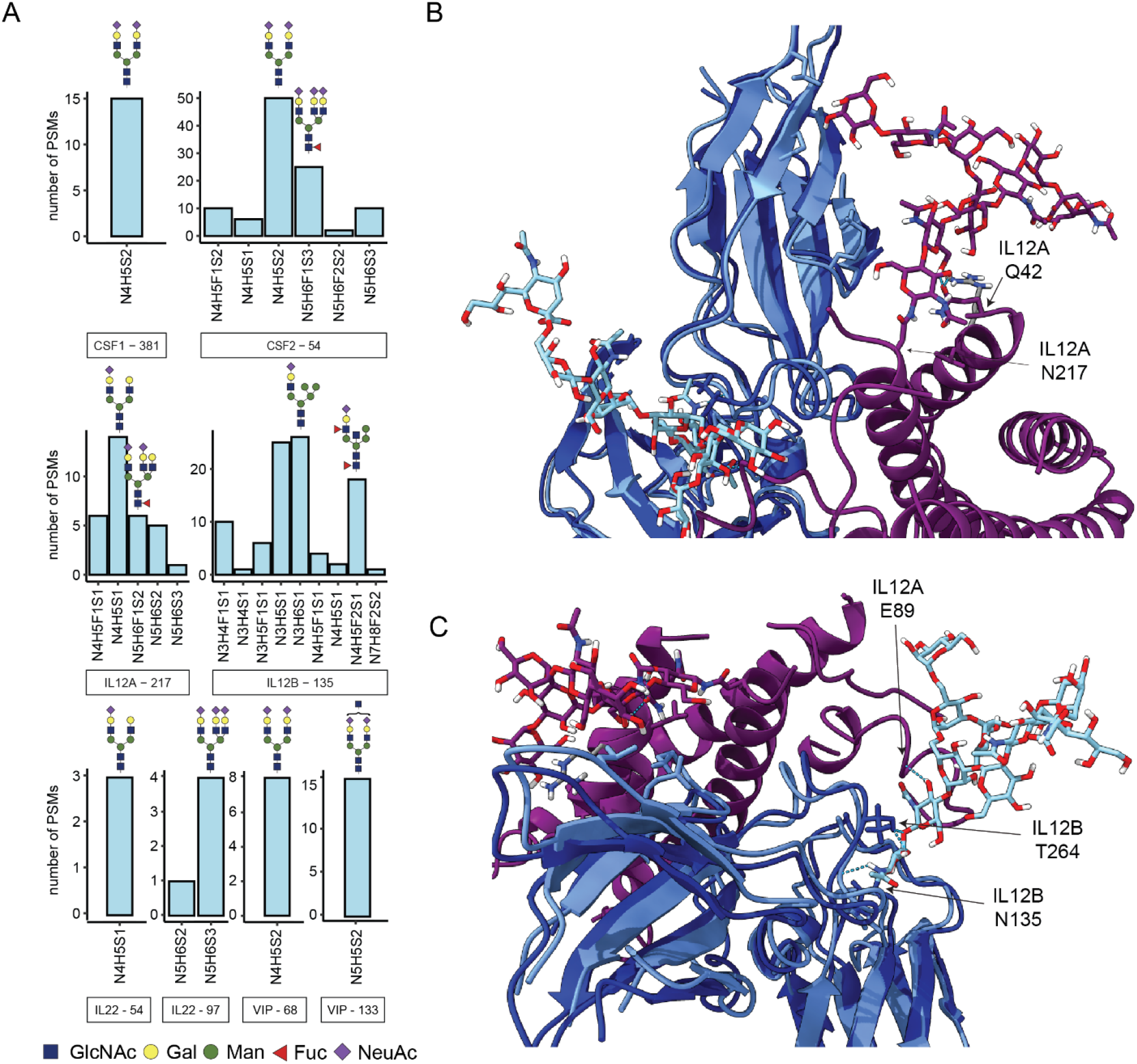
GlycoPSMs detected on low abundant cytokines in MS-based plasma proteomics. **A**) The number of PSMs depicts the total number of identified glyco-PSMs across all replicates and conditions in glycopeptide-enriched plasma. The protein name is indicated below the bar graph and the most abundant glycan structures are shown. **B and C**) X-ray structural model of IL-12, with the here observed most abundant glycans mapped onto these structures, where IL-12A and its glycan are purple, and IL-12B and its glycan are blue, with in light blue the protein structure as reported in the heterodimeric conformation (PDB 1F45), and in dark blue the overlayed monomeric structure (PDB 1F42). Light blue dashed lines represent possible hydrogen bonds between the glycans and proteins. The arrows indicate the glycosite and the interaction partners.

**Figure 6:**
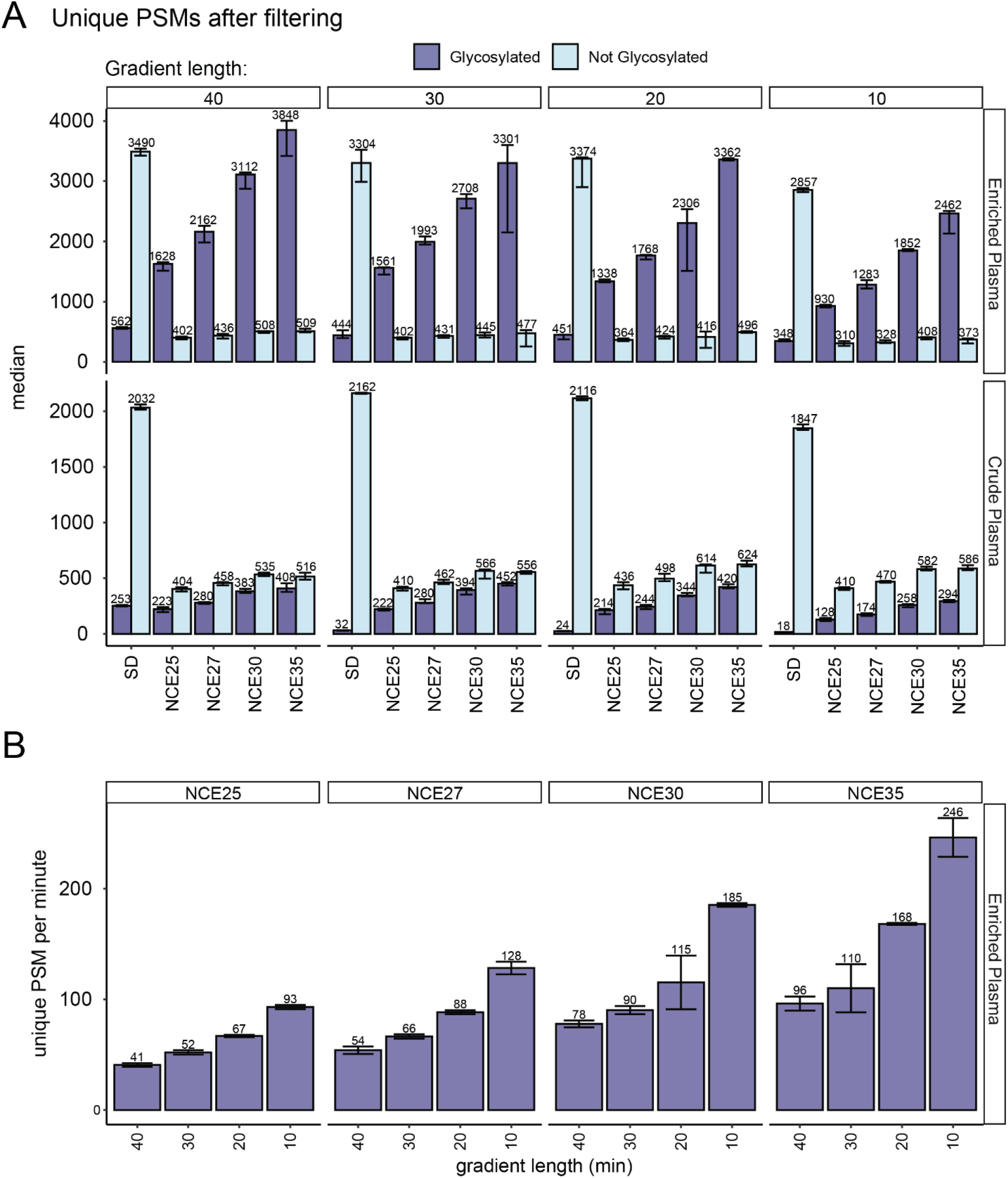
Evaluating the time-efficiency of the DIA glycoproteomics methods. **A**) Unique PSM count identified per method across four technical replicates in enriched plasma (top-panel) or crude plasma (bottom-panel). The bars illustrate the median values, while the error bars extend from the minimum to the maximum number identified out of the four replicates. **B**) Unique PSMs per minute in enriched plasma using different MS^2^ NCEs. The PSMs per minute is calculated as the median PSM count divided by gradient length. All data is filtered for 1% FDR and each unique PSM was identified in at least two out of four replicates for a single condition.

First, we focused on the interleukins. Interleukin (IL) 12 is a potent immunoregulatory cytokine crucial for the generation of cell-mediated immunity to intracellular pathogens^46^. It is a hetero-dimer, consisting of the sub-units IL-12A and IL-12B, of ∼25 and ∼37 kDa, respectively. For IL-12A we identify five different glycan compositions of complex type glycans on N217, a site that is however not annotated as a putative glycosite in UniProt. The detected glycans are N4H5S1, N4H5F1S1, N5H6S2, N5H6F1S2, and N5H6S3. Most of them seem to have one sialic acid less than the number of antennae. We were unable to establish whether the additional arm was branched or elongated. Manual analysis of the Y-ions in the different nGlycoDIA spectra did confirm that the fucose was most likely located on the core HexNAc. This N217 carries the *N*-glycan consensus motive and is located very near the C-terminus of IL-12A, (**<u>N</u>**AS), whereby Ser represents the C-terminal amino acid. We could trace no earlier structural or functional data on this intriguing C-terminal *N*-glycosylation.

For IL-12B we detected two distinct glycosylation sites, namely at N125 and N135. For site N125, we only detected a peptide with two amino acids, and although we detected two different glycans on it across multiple conditions, we felt this data is not sufficient to confidently assign the site. On site N135, which is annotated as putative glycosite in UniProt containing the consensus motif (**<u>N</u>**YS), we detected several hybrid type glycans (**Supplemental Figure S6**). These glycans contained at least one sialic acid and some were fucosylated with up to two fucose moieties as well. One of the glycoforms was consistently annotated as N4H5S2, however, after manual curation, we corrected this to N4H5F2S1 (**Supplemental Figure S6B**). This is a well-known mistake that occurs frequently when the annotation software picks the wrong peak during monoisotopic peak assignment. Another interesting feature on these hybrid glycans was the direct loss of an *N*-acetylhexosamine from the precursor upon fragmentation, suggesting the presence of a bisecting GlcNAc or additional antenna.

We mapped the sites and most abundant glycan structure found in our study for IL-12A and IL-12B on the reported X-ray structural models of the IL-12 heterodimer (PDB: 1F45) (**Figure 5B**)^48^. Only the site on IL-12A could be modeled this way, as the site for IL-12B was not in a solvent-accessible site in its configuration within the hetero-dimer. We were able to map the glycan on IL-12B on its monomeric configuration (PDB: 1F42) and overlaying it to the hetero-dimeric conformation (**Figure 5C**). Taking a closer look at the structure of the hetero-dimer and the glycans, we noticed that the core GlcNAcs from IL-12A-N217 formed hydrogen bonds with the sidechain of the Gln42 residue of IL-12A, which may aid in stabilizing its conformation upon hetero-dimerization (**Figure 5B**). The glycan core of IL-12B-N135 also formed intermolecular hydrogen bonds with the Thr264 sidechain of IL-12B and the second GlcNAc formed a hydrogen bond to the amide of the backbone at Glu89 of IL-12A, which is in a flexible region, possibly aiding in stabilization. For these X-ray studies, the protein was obtained by recombinant production in CHO cells and did contain glycans. In the structure of IL-12B another *N-*glycan was reported on IL-12B at site N222 (N200 by the author’s numbering), which we however did not detect in our study.

On IL-22, we experimentally detected two glycosylation sites, namely at N54 and N97. On both sites we only identified one glycan: N4H5S1 on site N54 (**<u>N</u>**RT), N5H6S3 on site N97 (**<u>N</u>**FT). Both these sites are annotated in NextProt as putative *N*-glycosylation sites. IL-22 is a 20 kDa α-helical cytokine that binds to a heterodimeric cell surface receptor composed of the IL-10R2 and IL-22R1 subunits. We mapped these two observed glycans on the structure of IL-22 when bound to the IL-22R1 receptor (PDB 3DLQ) (**Supplemental Figure S7A**)^49^. The glycan at N97 points away from the complex, while the glycan at N54 is pointing towards the receptor, especially with the glycan antenna. Previously, it has been reported that glycans or IL-22 (recombinantly produced in insect cells) did not interfere with the binding of IL-22 to its receptor and that the presence of absence of the glycans does not have a large impact on the conformation^50^. The glycans on these recombinant proteins were different from those detected here in human serum; the insect cell glycans were shorter and did not have antennae, while the here detected glycan is di-antennary and has a single negative charge on the sialic acid residue. Still, also in our model the glycan likely does not influence the binding to the receptor as it is not in proximity.

The macrophage colony-stimulating factor 1 (CSF1) promotes the release of pro-inflammatory chemokines, and thus plays an important role in innate immunity and inflammatory processes. This ∼60 kDa protein does carry more than a dozen *O*-glycosylation sites (not studied here) but has according to NextProt two putative *N*-glycosylation sites (N154 and N172, annotated by consensus sequence). Here, we did not find evidence for these two sites, albeit we confidently identified *N*-glycosylation at N381 (NHT), a site not annotated in NextProt. We found the site to be occupied with a N4H5S2 glycan, a di-antennary complex type glycan. There is an *O*-glycosylation site annotated in NextProt at the adjacent T383 site, located on the same tryptic peptide as N381, however the Y-ions series we identified supports that the annotation the *N-*glycan site at N381 is likely.

CSF2, also known as granulocyte-macrophage colony-stimulating factor, is a ∼15 kDa protein and has in NextProt also two putative *N*-glycosylation sites (N44 and N54), of which we here identified N54 to be occupied by various complex glycans, almost always fully sialylated and sometimes harboring also core fucosylation. For this cytokine at site N54 we therefore have multiple pieces of evidence that it can be detected in plasma (reported concentration in the Blood Atlas of 2.4x10^4^ pg/L) and that it is glycosylated.

We also detected the vasoactive intestinal peptide (VIP), which has no annotated glycosites in NextProt and a reported concentration of 1.2x10^5^ pg/L in the Blood Atlas. On VIP we identified two *N*-glycosylation sites: N68 (**<u>N</u>**DT) and N133 (**N**YT). The former is located on the propeptide and the latter is located on the vasoactive intestinal peptide. Here we detected glycopeptides whereby N68 was occupied by an N4H5S2, while N133 was found to be occupied with N5H5S2. Recently, a cryoEM structure was reported revealing the interaction of a VIP peptide (125-152) coupled to its G protein-coupled pituitary adenylate cyclase-activating polypeptide (PACAP) type 1 receptor (PAC1R). The authors compared the binding between PAC1R and VIP to the binding of PAC1R to PACAP, the latter having a 1000-fold higher affinity while sharing nearly 70% sequence identity^51,52^. Interestingly, one of the sequence differences between VIP and PACAP is in the *N*-glycan consensus motif, although the binding between PAC1R and VIP was only investigated using a non-glycosylated VIP peptide. The VIP peptide agonist was shown to bind in the relatively large *N*-terminal extracellular domain (ECD), facilitating engagement of the peptide N-terminal activation domain with the extracellular loops (ECLs) and transmembrane helices (TMs) of the receptor core. We modeled the here observed glycan on this structure (PDB 8E3Z) (**Supplemental Figure 6B**). As the peptide is fully situated within the ECD domain, with the glycan sticking out, the here reported glycan moiety very likely affects this binding.

Although they are normally extremely low abundant, we detected several more cytokines exclusively by our nGlycoDIA approach, and only in the glycopeptide enriched plasma, albeit with less evidence. These included IL4 (N47 and N114), IL34 (N77), IL24 (N127), FMS-like tyrosine kinase 3 ligand (FLT3LG) and tumor necrosis factor superfamily member 11 (TNFSF11). The fact that we detect all these cytokines was rather remarkable as most of them have concentrations in the order or 10^4^-10^5^ pg/L in plasma of healthy donors (as analyzed here), whereas the concentration of, for instance, the immunoglobulins and some of the serpins are 10^10^-10^11^ pg/L in plasma. This demonstrates an apparent dynamic range of 10^6^ in glycoprotein abundance using our nGlycoDIA approach on enriched plasma.

### Towards high-throughput and high-sensitivity plasma glycoproteomics

Although 40 min gradients are already relatively short for glycoproteomics, to investigate larger clinical cohorts of plasma samples for glycoprotein biomarkers any reduction in analysis time would be beneficial. With the optimized nGlycoDIA method in hand, an effort was made to decrease the time of each run. We tested the optimized nGlycoDIA method with different gradient lengths (40 min, 30 min, 20 min and 10 min). In fairness, note that these numbers represent the LC gradient length, and not the total time of the LC method as the latter includes an additional 6-minute column wash.

In the glycopeptide enriched plasma, we could detect a decrease in the number of unique glycopeptide PSMs with shorter gradient times. Still, just about 10% of all unique PSMs were lost when the gradient time was reduced from 40 to 20 minutes, while ∼40% was lost when decreasing the gradient time to 10 minutes (**Figure 6A**). The number of annotated PSMs without glycan decreased at maximum 25% from 40 to 10 minutes when using nGlycoDIA in enriched plasma. For the crude plasma, our data indicated that extending the gradient past 20 minutes did not lead to an increase of unique glycoPSMs. The number of proteins and glycosites revealed similar trends. Another way to examine the time-efficiency is to look at the unique identifications per minute (**Figure 6B**). Consistently for all methods, the glycosylated identifications per minute decreased with gradient duration, the 10-minute gradient being still the most efficient. As far as we know these represent some of the most data-rich plasma glycoproteomics data with such short (10 min) gradients.

When comparing the different gradients and different NCEs, we again observed a huge overlap (∼80%) between identified PSMs, similar to the overlap between replicates as seen in the 40-minute measurements (**Supplemental Figure S3A and S3B**). When clustering all the data, the first separation was based on enriched vs. non-enriched plasma (**Supplemental Figure S8**). Subsequent grouping was primarily based on the number of identifications in each dataset, further demonstrating the reproducibility of the nGlycoDIA approach.

Lastly, we compared the dynamic range between the 10 minutes and 40 minutes gradients. Here we saw that in the 10 minutes approach, compared to 40 minutes, we detected fewer low-concentration glycoproteins (**Supplemental Figure S9A**), *i*.*e*., 32% fewer proteins in the µg/L range and almost 50% fewer proteins in the ng/L range. Additionally, there was a decrease in annotatable microheterogeneity in the µg/L range, albeit not in the other concentration ranges (**Supplemental Figure S9B**).

## Discussion

In this study, we evaluated the novel Orbitrap Astral MS instrument for plasma glycoproteomics, demonstrating the potential of narrow-window data-independent acquisition (nDIA) for analyzing glycopeptides (nGlycoDIA). With this method, using crude plasma, we were able to annotate around 400 unique glycosylated PSMs per run, covering 60 glycoproteins and 186 glycosites. Remarkably, without enrichment, the 20-minute method was already sufficient to maximize detection of glycopeptides in this sample. However, following a simple and cost-effective cotton based HILIC glycopeptide enrichment, nGlycoDIA resulted in the identification of 3000 unique glycosylated PSMs using 40 minute LC gradients, covering in total 181 glycoproteins and 436 glycosites. These 181 proteins spanned a dynamic range across 7 orders of magnitude, significantly surpassing previous studies which applied glycopeptide enrichments on.plasma ^13,26,43^

This is as far as we are aware one of the first application of DIA for plasma glycoproteomics and provides a substantial advance over previous methods used in other and our laboratories. This advance can be largely contributed to the higher speed of the Astral analyzer, but also the use of for glycopeptides optimized narrow-window DIA, allowing the data to be analyzed by a DDA-like computational processing workflow. DIA is for glycoproteomics much more challenging than for ‘standard’ proteomics, due to many factors: amongst them are the fragmentation efficiency of glycopeptides and the high abundance of oxonium ions that are not specific to unique glycopeptides. As this is the first nGlycoDIA method for plasma glycoproteomics, and one of the first test of the Astral mass analyzer for glycoproteomics, it is hard to compare our data with previous studies.

Recently, a plasma proteomics study was published using narrow-window DIA on the Orbitrap Astral MS^29^. Although this study did not target glycopeptides and used a lower MS^1^ precursor range (380-980 *m/z*), we were interested in comparing the proteins identified in that study to those identified using nGlycoDIA. At first glance, the here reported number of 181 detected glycoproteins may seem rather low, as in the previous mentioned study more than 4500 proteins were identified. However, it should be noted that the latter study used an extracellular vesicle (EV)-based enrichment to reduce the dynamic range of the plasma sample. In that same study, 667 proteins were identified from “crude plasma” without EV-based enrichment, which is still about 3-fold more than we report. One would expect *a priori* that all 181 glycoproteins we identified here would also have been detected in this much “deeper” study, that used the narrow-window DIA method^29^. However, out of these 181 just 43 (24%) were also identified by Heil *et al*. in the crude plasma. Even in their dataset after EV-based enrichment (4500 identification) only 126 (70%) of the glycoproteins we identified are present, leaving 55 glycoproteins uniquely identified by the here described dedicated nGlycoDIA approach (30%). These 55 are almost all low abundant glycoproteins which have been described as genuine plasma proteins, often only detected by targeted highly sensitive enzyme-linked immunosorbent (*ELISA*) assays^35^. Their reported plasma concentrations range from 1 mg/L down to 1 ng/L. For most of these low abundant proteins, glycosylation has not been experimentally reported by MS-based plasma proteomics, although many of the here observed glycosites are annotated as putative *N*-glycosites in the public depositories UniProt or NextProt. Evidently, although ELISA assays can be very sensitive, they are not directly useful in detecting PTMs. Amongst these low abundant proteins were several glycosylated cytokines. This data will be valuable for understanding the role of glycosylation in the secretion and function of these cytokines.

One initial conclusion we can draw is that, although the absolute number of identified proteins in our nGlycoDIA proteomics approach is only a fraction of the ∼4500 proteins reported in the recent nDIA EV-enriched plasma studies on the same Orbitrap Astral system, we were able to expose a unique part of the plasma proteome. The two-fold enrichment (both offline using HILIC-based glycopeptide enrichment and in the gas-phase with the optimized MS^1^ window) and the speed and sensitivity of the Orbitrap Astral MS and the newly developed nGlycoDIA all aided to the expanded view of plasma glycoproteomics and the coverage of a broad dynamic range (10^7^).

However, our study also encountered limitations. In our data-analysis we were limited in our choice of analysis software, since DIA, especially library-free DIA, is not yet well established for glycoproteomics. At the time of analysis Byonic was the only analysis software available that accepted DIA data format and was able to assign the MS1 precursor masses and charge states. Data quality might have improved with the use of broader windows, which would allow more time per scan. Recently, a similar Byonic strategy was used on DIA data with 16 *m/z* windows^27^. However, this was performed on a purified protein, thus the data was less complex, and the chance of chimeric spectra is lower than when plasma is used. Therefore, we decided to use very narrow windows (3 Th) to reduce the possible occurrence of chimeric spectra. If software becomes available to analyze library-free DIA data, we could potentially enhance data quality further by utilizing broader windows and allowing more time per scan. Furthermore, we used four different collision energies here, which each provided important information for compositional and structural annotation of the glycopeptides. Being able to use stepping collision energy without sacrificing speed would enable all complementary information to be acquired in a single scan, as is often used in DDA glycoproteomics^14,15,54^. However, applying stepping HCD in this case meant we had to either reduce the cycle time, the injection time, or the MS1 scan range, or we had to increase the window size which would lead to more chimeric spectra. To stay as close to DDA type data and reduce chimeric spectra, we chose to work with a single collision energy for each run. Moreover, manual analysis remained crucial for detailed glycan structure identification, as search engines only provide compositional information, and are still prone to incorrect structural annotations.

Despite these limitations, the data produced by the Orbitrap Astral MS facilitated in-depth analysis of a large set of proteins in plasma. We showed that qualitative data independent acquisition from a single plasma sample can provide rich information on glycopeptide diversity. This aligns with our primary objective and underscores the potential for future research, exploring even more extensive glycoproteomic signatures and glycan structures on low abundant proteins.

Future studies could aim to address quantification challenges, perhaps by integrating different software algorithms. Furthermore, the ability to analyze glycan signatures on very low abundant proteins without immunoprecipitation or fractionation, opens the door for novel insights in disease mechanisms. Additionally, this method, and especially the ability to do in-depth glycoproteomics in only 10 minutes per sample, opens pathways for broader applications, including clinical screenings, high-throughput analysis and biomedical research, where insights into glycan structures could inform disease diagnosis or treatment strategies. As a by-product of our work, we hypothesize that quantitative analysis of very low abundant cytokines in plasma may be helped by using glycopeptide enrichment, as several of the here detected cytokines are typically not observed in standard plasma proteomics.

Overall, this work establishes a good foundation for in-depth plasma glycoproteomics, where a simple and cost-effective enrichment followed by a short DIA method enables the identification of a broad range of glycoproteins and their glycans.

## Acknowledgements

This research received funding through the Dutch Research Council (NWO) funding the X-omics Road Map program (project 184.034.019). AJRH acknowledges support from NWO through the Spinoza Award SPI.2017.028.

## Author Contributions

SJ, KRR, AAM and AJRH conceptualized the project; SJ, MZ, and AP performed the experiments; SJ, MZ, AP, and ED worked on method development; SJ and DS performed data analysis and visualization; AAM and AJRH provided resources; SJ and AJRH were responsible for the writing; all authors further edited the work and critically revised the final version of the manuscript.

## Disclosure Declaration of competing

### interest

The authors declare the following potential competing interests: MZ, AP, ED and AAM are employees of Thermo Fisher Scientific, the manufacturer of the mass spectrometers used in this study.

## Data Availability

All raw data files and Byonic output files used in this study has been reposited to the MASSive repository, and will be made available upon request.

