## Supplementary Figures S1 to S9 for "Narrow window data-independent acquisition on the Orbitrap Astral Mass Spectrometer enables fast and deep coverage of the plasma glycoproteome"

### Index


| | |
|---|---|
| Supplemental Figure S1 | 2 |
| Supplemental Figure S2 | 3 |
| Supplemental Figure S3 | 4 |
| Supplemental Figure S4 | 6 |
| Supplemental Figure S5 | 7 |
| Supplemental Figure S6 | 8 |
| Supplemental Figure S7 | 9 |
| Supplemental Figure S8 | 10 |
| Supplemental Figure S9 | 11 |

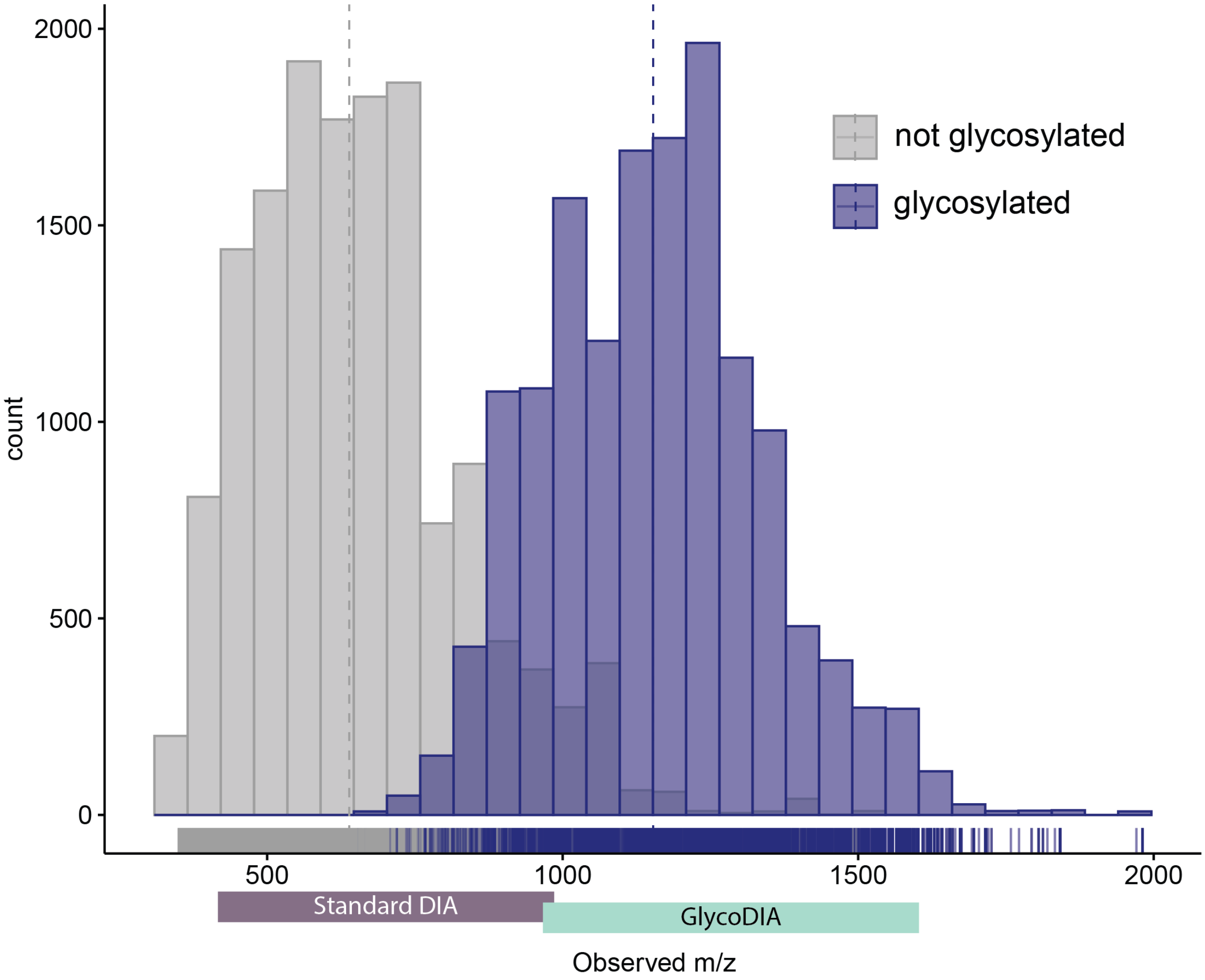


**Supplemental Figure S1: Distributionof precursor *m/z* of non-glycosylated and glycosylated peptides, used for MS1 scan range optimization.** The glycopeptide enriched plasma was analyzed on an Exploris 480 and a histogram was made to explore the precursor *m/z* values of glycosylated (dark blue) and non-glycosylated (grey) peptides. Here, we find that glycosylated peptides have a distinct *m/z* precursor distribution, which is rather different from non-modified peptides. Comparing this to the range used in the recent narrow-window DIA glycoproteomics workflows (annotated as Standard DIA), we see that only a marginal fraction of glycopeptides will be targeted with this earlier method. Therefore, we moved here the $MS^1$ scan range up to between 950-1655 *m/z*, as this range will cover most glycopeptide precursors, Additionally, we expect this to result in a orthogonal degree of glycopeptide enrichment, i.e. in the gas phase.

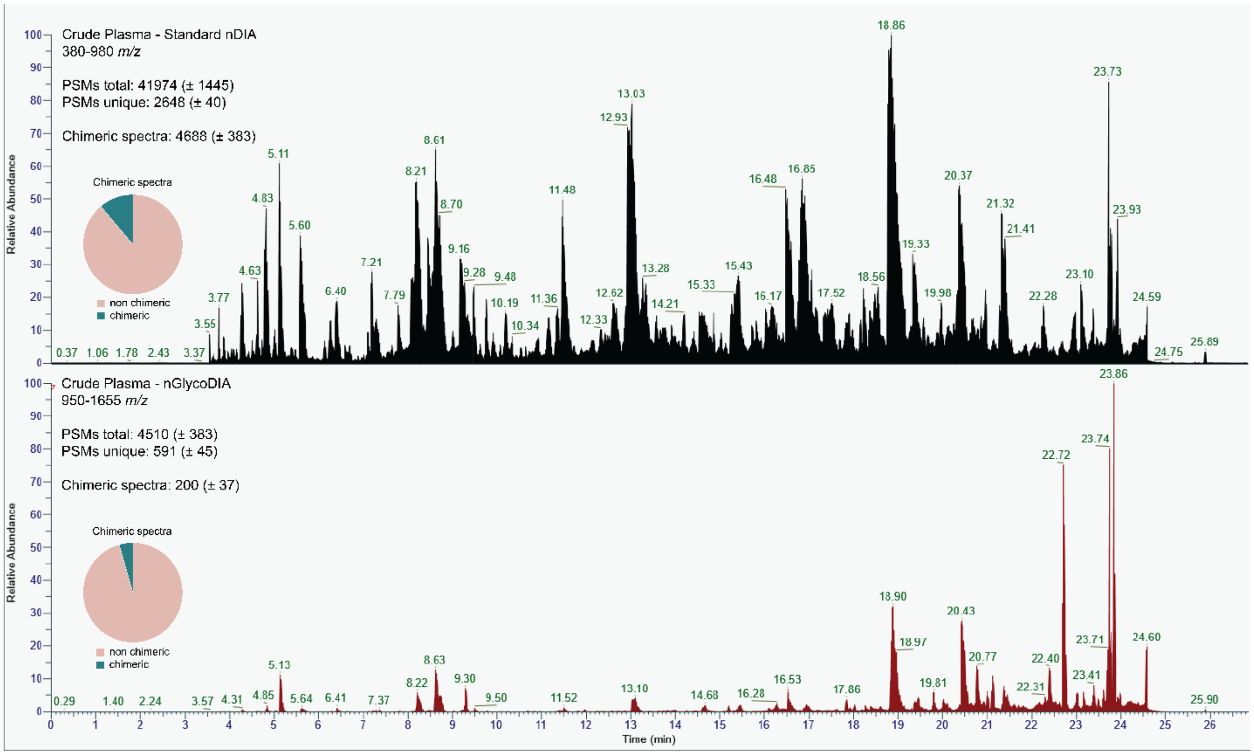


**Supplemental Figure S2: Consequences of shifting the mass window for proteomics.** Depicted are the total ion current (TIC) chromatograms of an analysis of crude plasma using the standard nDIA method (top) and the nGlycoDIA method (bottom). The intensity is plotted as the relative abundance, and the maximum intensities of both analyses are very comparable, 3.99E10 and 3.30E10 for the top and bottom analysis, respectively. Metrics given are the avarage number of total PSMs, unique PSMs, and chimeric spectra, with between brackets the standard deviation. From these metrics it is evident that nGlycoDIA leads to a 9-fold reduction in PSMs, and a 5-fold reduction of unique PSMs, but also to a more than 20-fold reduction of chimeric spectra. To furhter illustrate that the percentage of chimeric spectra is lower in nGlycoDIA, a piechart is shown of non-chimeric (pink) and chimeric (blue) spectra.

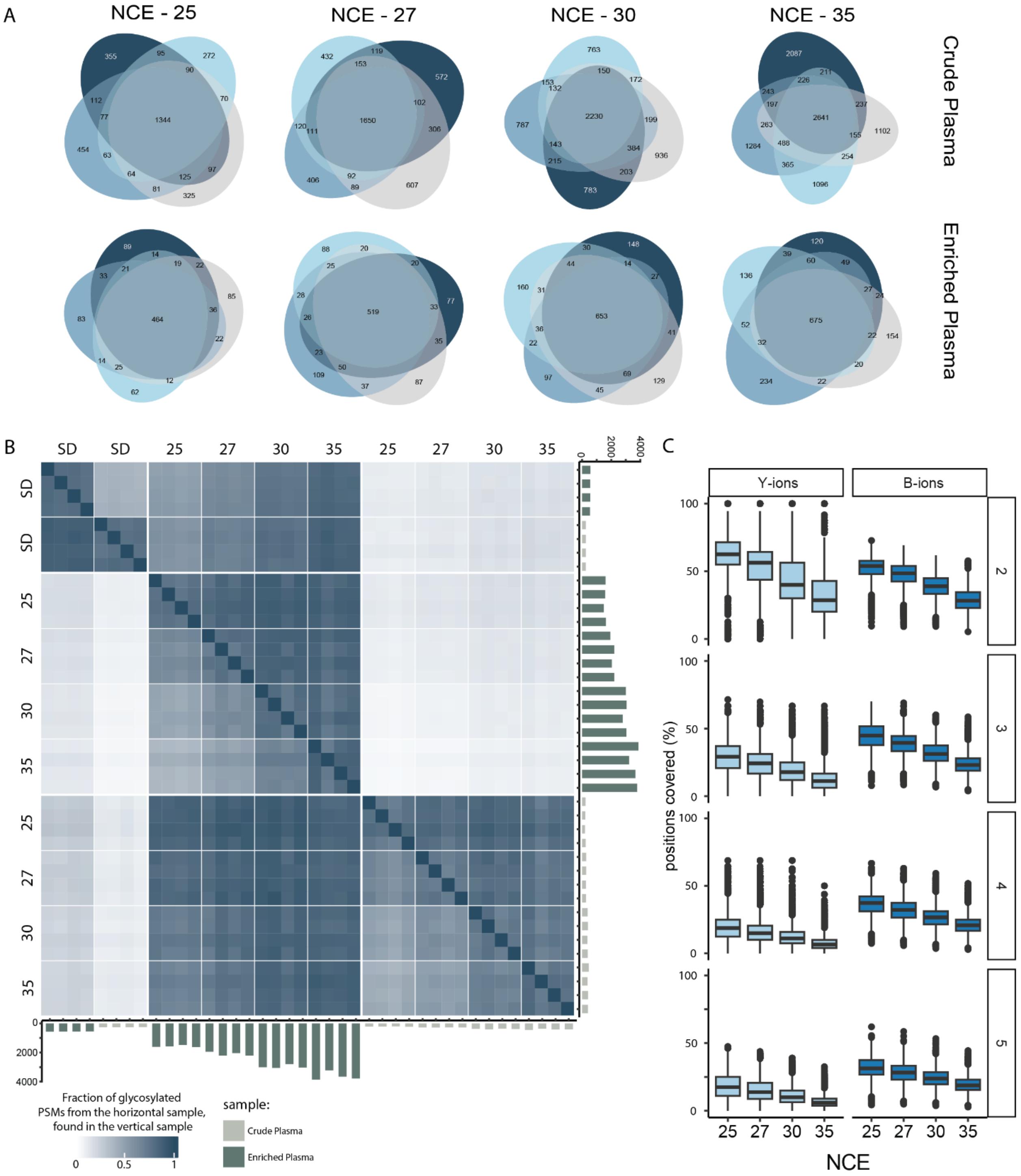


**Supplemental Figure 3: Exploring the quality and reproducibility of the nGlycoDIA method. A)** Venn diagrams of four replicates for each collision energy (NCE) used before applying our filtering criterium of a PSM being annotated in at least 2 out of 4 replicates in at least 1 condition. Evidently, when a PSM is annotated in at least 2 replicates, it is usually annotated in all replicates; illustrating the reproducibility of the method. **B)** Heatmap of the overlap in detected glycopeptides between samples, where the color indicates the number of glycopeptides (unique glycan + modifications + peptide) from the horizontal sample that was also identified in the vertical sample. On both axis barplots depict the number of unique glycan PSMs that were found in each individual sample, the colors of the bars refer to which sample was used: light green for crude plasma, and dark green for enriched plasma. This heatmap displays the big overlap between replicates and conditions. Moreover, smaller datasets with less unique glycoPSMs are generally fully contained within the larger datasets, indicating that the increase in unique glycoPSMs is an enrichment and not an artifact. Values on the x- and y-axis refer to the MS/MS method used, where SD refers to the standard DIA method, and 25, 27, 30, and 35 are nGlycoDIA methods with the respective value as NCE. **C)** Boxplots illustrating the percentage of theoretical positions covered of glycan-specific ions (Y-ions and B/oxonium-ions). The number of theoretical positions is calculated by matching the glycan composition to available structures in the GNOme database (https://pages.glycosmos.org/gnome/public/StructureBrowser.html), calculating all possible fragments in all possible charge states (with the maximum charge state being the precursor charge state), and matching these to the peaks in the data with an 20 ppm mass error. Because the number of positions increase with the precursor charge state, the 4 most abundant charge states are depicted here and separated in different panels. All charge states demonstrate the same phenomenon, both Y- and B-ion coverage decreases as the used NCE increases.

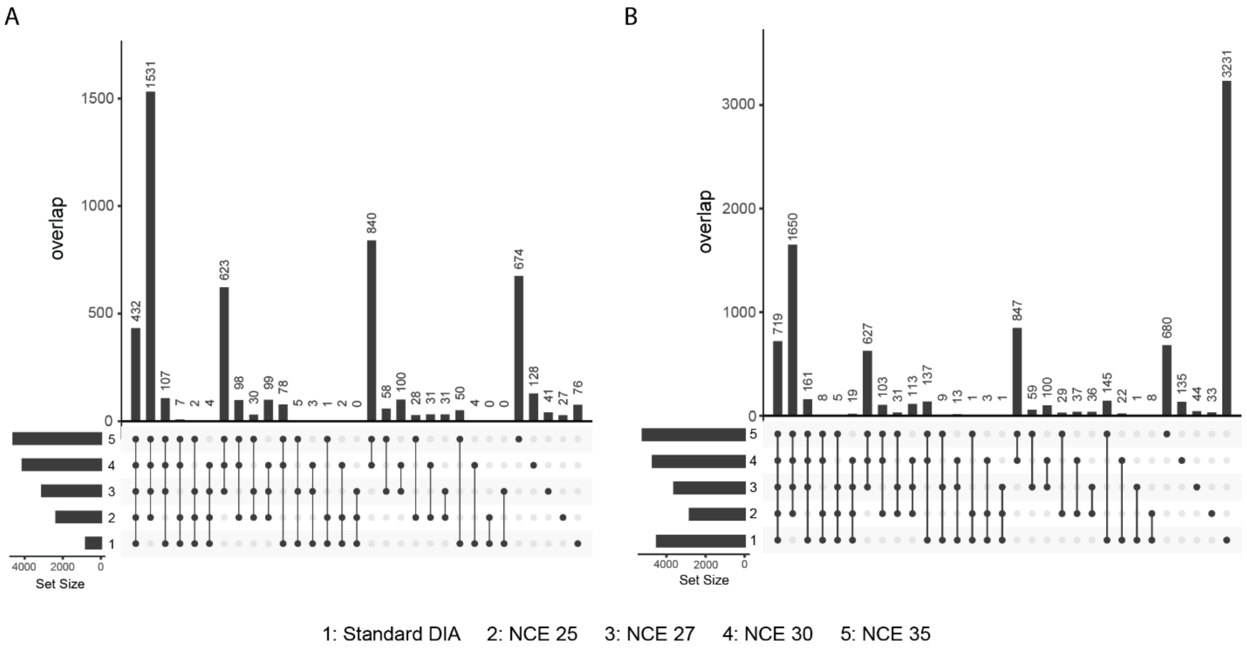


**Supplemental Figure S4: Upset plots depicting the overlap in detected glycopeptides between the different DIA and nGlycoDIA MS methods.** Upset plots of the overlap in **A)** unique glyco PSMs and **B)** all unique PSMs. Data was filtered for 1% FDR and, to be included, each unique PSMS had to be identified in at least 2 out of 4 replicates for a single condition. De total size of the dataset is displayed in the bar graph in the bottom left corner, and samples are annotated as numbers 1 to 5, where 1 is Standard DIA, 2 is nGlycoDIA with NCE 25%, 3 is nGlycoDIA with NCE 27%, 4 is nGlycoDIA with NCE 30%, and 5 is nGlycoDIA with NCE 35%. The same data is displayed as a Venn Diagram in **Figure 2D and 2E** of the main text.

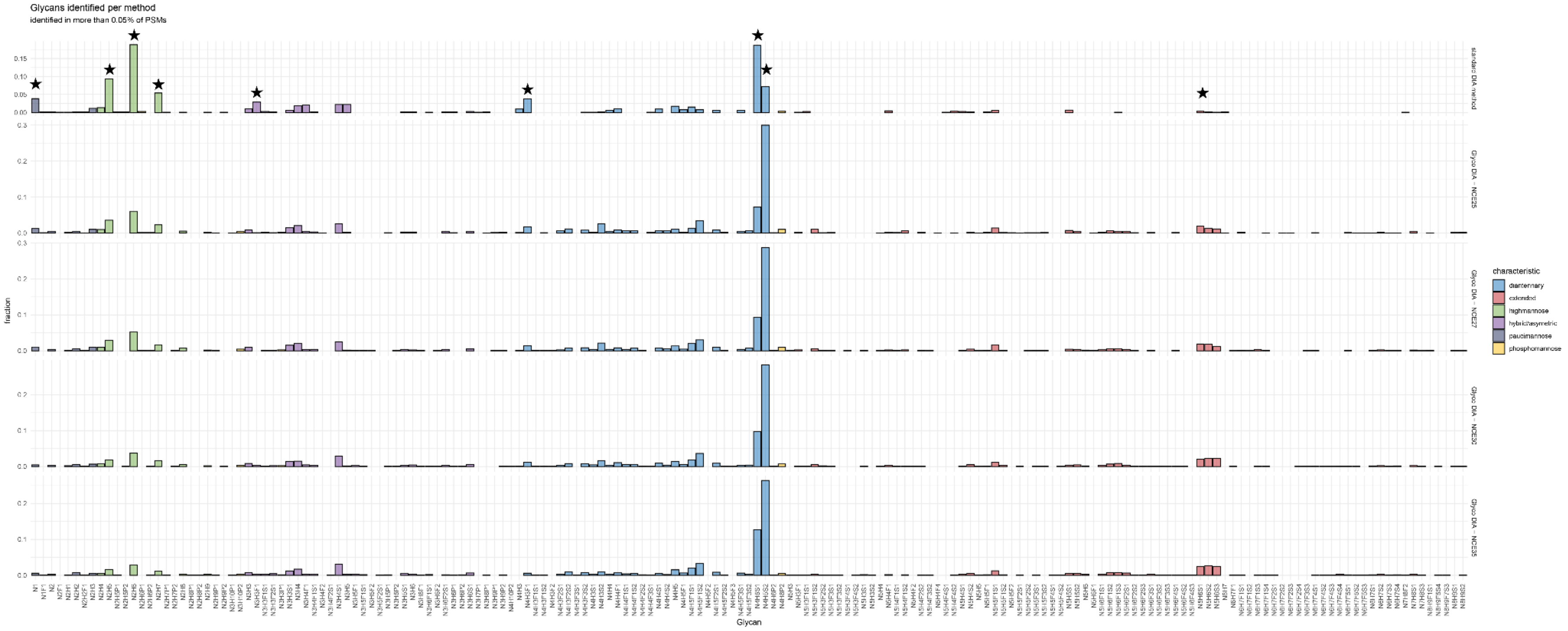


**Supplemental Figure S5: Cumulative distribution of detected glycans per MS/MS method.** Plotted here is the fraction of the total glycosylated PSMs bearing a specific glycan composition as annotated on the x-axis. The data is filtered for a minimum abundance of 0.05% (or fraction of 0.0005). From top to bottom, the methods are standard DIA, nGlycoDIA NCE25, nGlycoDIA NCE27, nGlycoDIA NCE30, and GlycoDA NCE35. The latter four are almost identical, while the standard DIA is quite different, with the most clearly different peaks annotated with the stars. We see that there are more paucimannose glycans, and more high-mannose in the standard DIA approach, compared to the di-antennary glycans that are the most abundant in all other methods. Additionally, the most found glycan in nGlycoDIA (N4H5S2, almost found on 30% on all PSMs) is only found on 5% of glycoPSMs in the standard DIA method. This figure clearly demonstrates that the standard DIA method created a bias in the detected glycopeptides, while collision energy itself did not introduce glycan bias. Evidently, peptides carrying paucimannose glycans are smaller than when harboring more complex glycans structure and may this better fall within the lower *m/z* window applied in the standard DIA method.

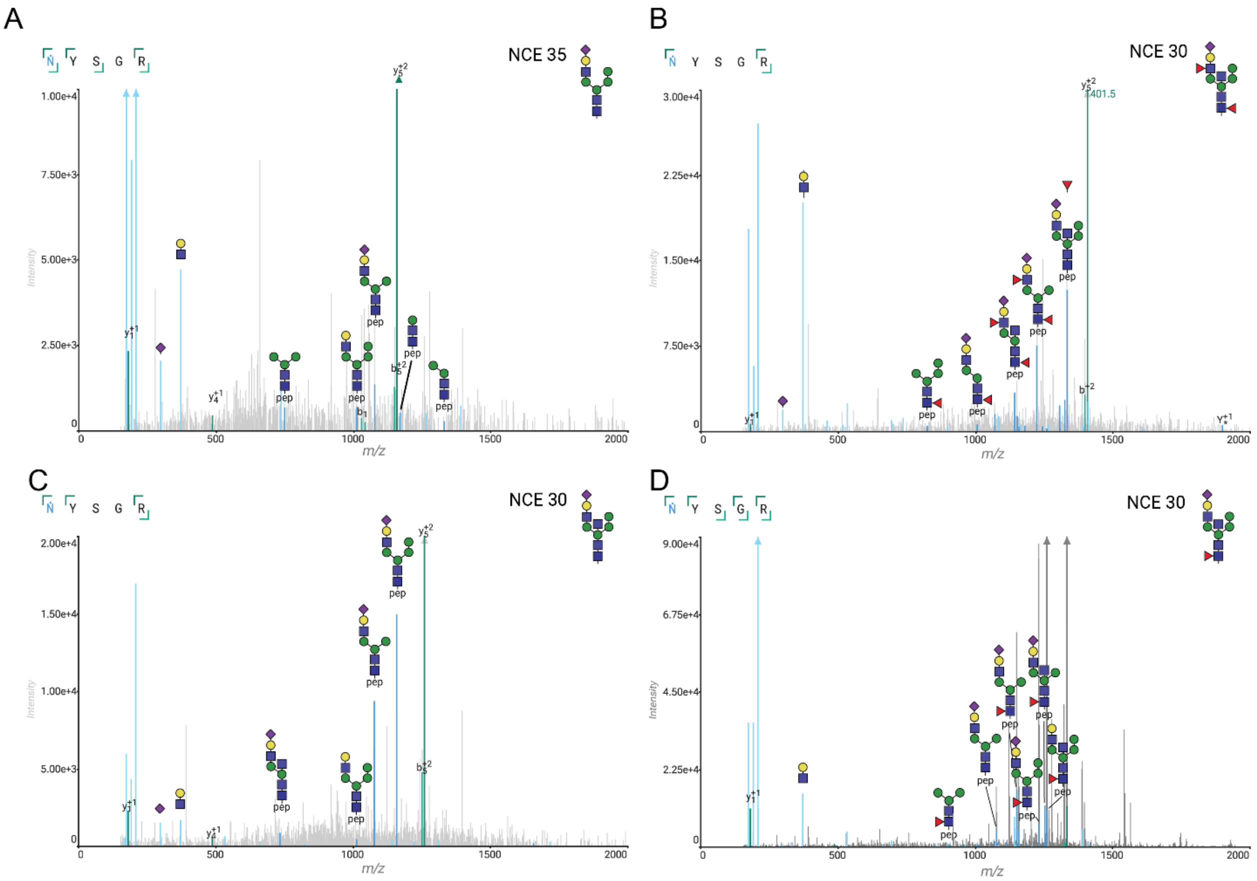


**Supplemental Figure S6: annotated spectra visualizing the glycans detected on IL12B-N135.** Annotated spectra of different glycans at different collision energies as depicted in the right top corner, next to the assigned glycan structure. The left top corner shows the peptide sequence with the fragment coverage of b- and y- ions. Glycan B- and Y-ions are annotated in light and dark blue, respectively. Most abundant Y- and oxonium ions are annotated with their proposed fragments. The glycan in panel **B** was originally annotated as N4H5S2, however, the Y-ions clearly show the loss of a fucose (mass loss of 146 Da) as its most abundant fragment. There are also fragments indicative of the hybrid structure and core fucosylation (the pep+N2H4F1 fragment) and we see fragments suggesting bisection, as also seen in panel **C** and **D**. This annotation, however, does not exclude other configurations or that the spectra are chimeric.

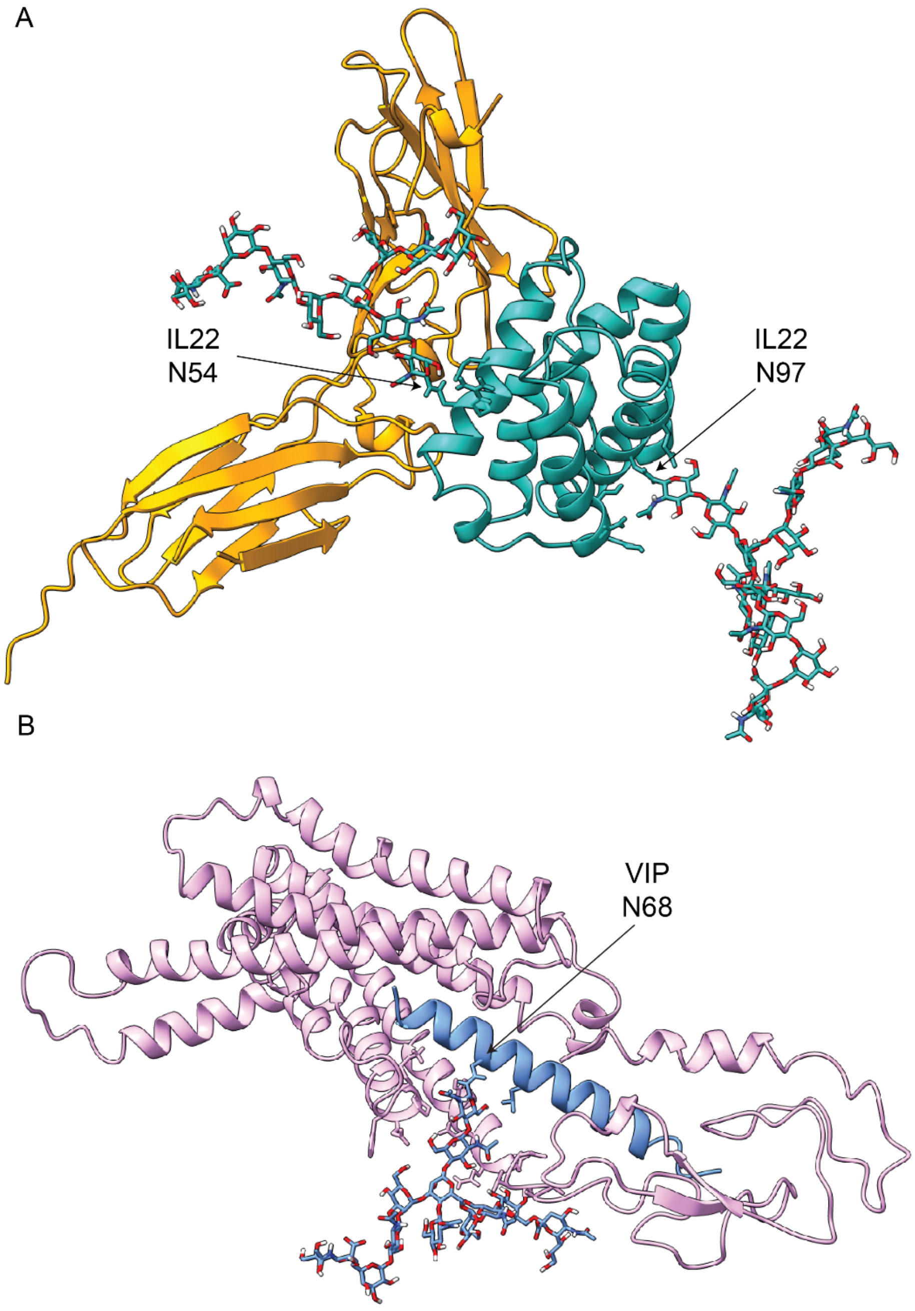


**Supplemental Figure S7: Structural models of the cytokines IL22 and VIP interacting with their receptor, with modeled in the here observed glycans. A**) The identified glycans are mapped on the crystal structure (PDB 3DLQ) of IL22 (turquoise) and its receptor (yellow). The glycan on N97 points away from the interaction interface, and seems not to interfere with receptor binding, while the glycan on N54 may point more towards the receptor. **B**) The identified glycan is mapped on the cryo-EM structure of VIP (blue) when bound to its receptor PAC1R (pink) (PDB 8E3Z). VIP is buried inside the extracellular domain of PAC1R, and thus this glycan may influence the binding of VIP to its receptor. Notably, the structure was resolved using non-glycosylated variants.

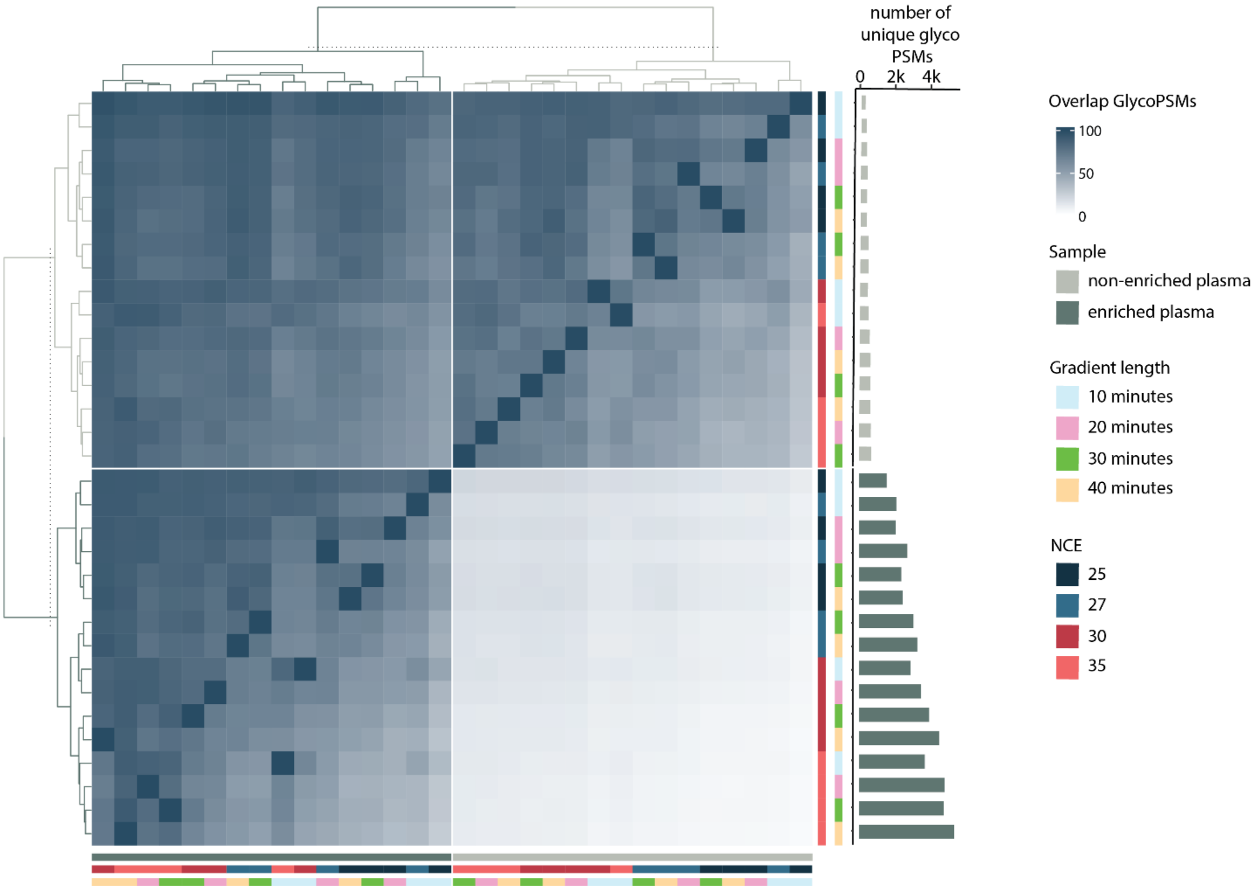


**Supplemental Figure S8**: **Overlap between identified glycosylated peptides using different gradient lengths and collision energies**. The heatmap color indicates the percentage of glycopeptides identified in the horizontal sample relative to the vertical sample. Sample identity is indicated by colored bars on the right and bottom sides. Samples primarily clustered based on sample identity (*e.g.* enriched or crude plasma), as well as on the total number of identified unique PSMs, as shown on the right axis.

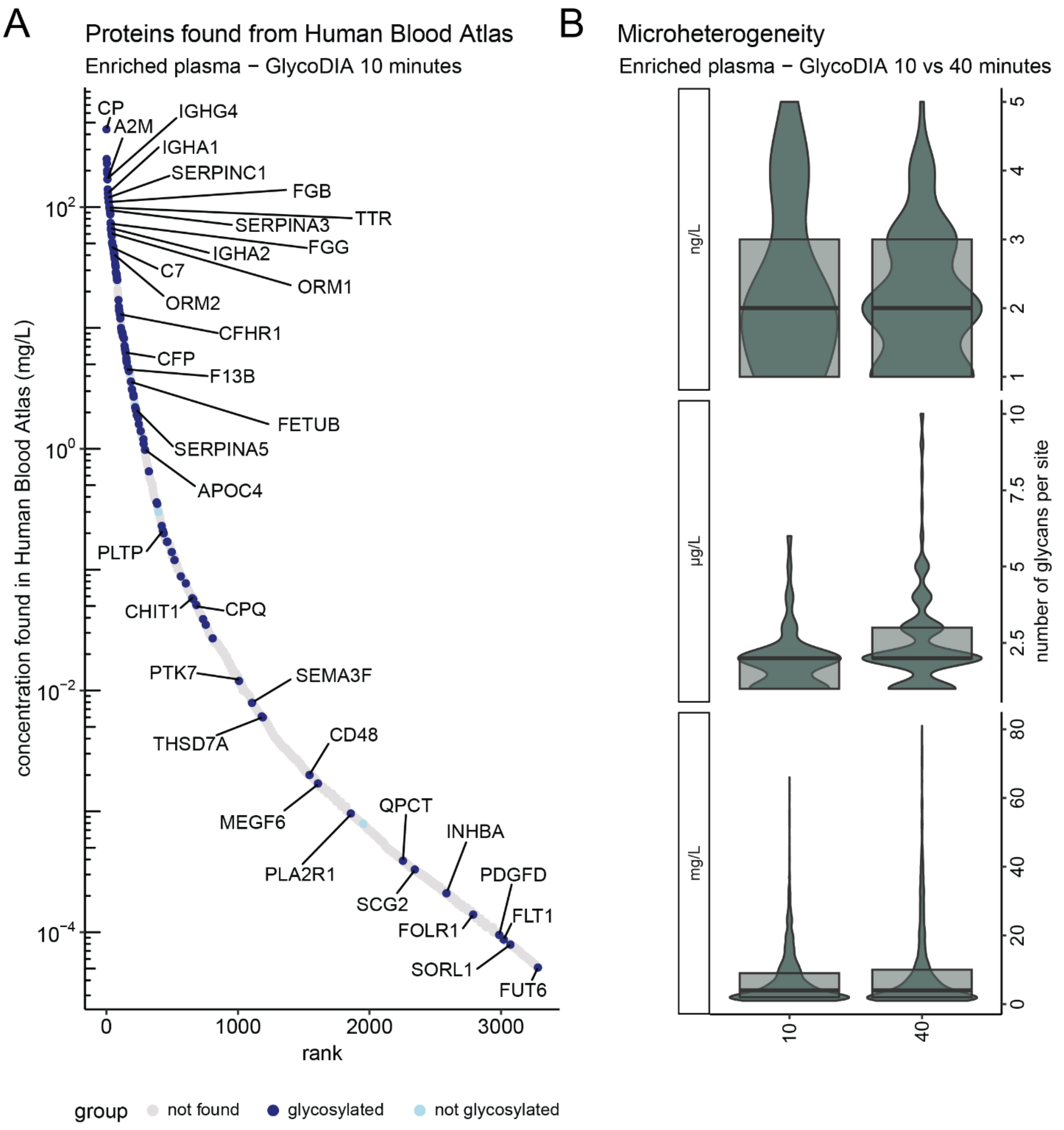


**Supplemental Figure S9: Comparison of depth and microheterogeneity as observed when using a 10- or 40-minute LC-MS gradient using nGlycoDIA on a glycopeptide enriched plasma sample**. **A**) The concentration of proteins identified compared to the reported concentrations as provided in the Human Blood Atlas. Even when using the 10 min gradient still a dynamic range of $10^6$ is covered **B**) Violin plot of number of unique glycans identified per site. Data is grouped on the reported abundance of the proteins in the Human Blood Atlas, where mg/L ranges between 999 - 1 mg/L, µg/L between 999 - 1 µg/L, and ng/L below 999 ng/L. The overlaying boxplot show the median value, and the values of the upper and lower quartile. Note that the scales of the y-axes are different.
